# An ultra-conserved ARF-DNA interface underlies auxin-triggered transcriptional response

**DOI:** 10.1101/2024.10.31.621286

**Authors:** Juriaan Rienstra, Polet Vanessa Carrillo Carrasco, Jorge Hernandez Garcia, Dolf Weijers

## Abstract

Auxin Response Factor (ARF) plant transcription factors are the key effectors in auxin signalling. Their DNA-Binding Domain (DBD) contains a B3 domain that allows base-specific interactions with Auxin Response Elements (AuxREs) in DNA target sites. Land plants encode three phylogenetically distinct ARF classes: the closely related A- and B-classes have overlapping DNA binding properties, contrasting with the different DNA-binding properties of the divergent C-class ARFs. ARF DNA-binding divergence likely occurred early in the evolution of the gene family, but the molecular determinants underlying it remain unclear. Here, we show that the B3 DNA-binding residues are deeply conserved in ARFs, and variability within these is only present in tracheophytes, correlating with greatly expanded ARF families. Using the liverwort *Marchantia polymorpha*, we confirm the essential role of conserved DNA-contacting residues for ARF function. We further show that ARF B3-AuxRE interfaces are not mutation-tolerant, suggesting low evolvability that has led to the ultra-conservation of the B3-DNA interface between ARF classes. Our data support the almost complete interchangeability between A/B-class ARF B3 by performing interspecies domain swaps, even between lineages that diverged over half a billion years ago. Our analysis further suggests that DNA-binding specificity diverged early during ARF evolution in a common streptophyte ancestor, followed by strong selection as part of a competition-based auxin response system

**Significance Statement:** Auxin response evolved nearly half a billion years ago in the earliest land plants. Auxin response is mediated by a family of DNA-binding ARF transcription factors. It has been unclear if and how the ARF family has evolved. In this paper, the authors show that the central protein-DNA interface that defines the genes that are under auxin control has remained essentially unchanged throughout auxin response evolution, explaining how auxin has become a dominant signal controlling growth and development in all land plants.

## Introduction

The plant signalling molecule auxin coordinates various developmental programs (1-3). This process is mainly governed by the nuclear auxin pathway (NAP), which is composed of three dedicated protein families: the TIR1/AFB receptors, the Aux/IAA co-repressors, and the final effectors, the Auxin Response Factor (ARF) family of DNA-binding transcription factors (3-7). Under low auxin conditions, Aux/IAA bind to and inhibit ARF function. When auxin levels increase, auxin triggers the formation of a receptor and co-repressor complex, destabilizing the latter. This liberates the ARFs to perform their intrinsic transcriptional function. Genes are auxin-dependent when there are specific cis-regulatory elements in their promoter: the Auxin Response Elements (AuxRE)(8-10).

ARF architecture shows three major and independent domains: the DNA-binding domain (DBD), the middle region (MR) and the Phox and Bhem 1 (PB1) domain. The MR defines whether ARFs transcriptionally activate or repress, while the PB1 domain drives ARF homotypic oligomerization, and heterotypic interaction with the Aux/IAA repressors (11). The PB1 domain also indirectly affects DNA-binding affinity through its role in cooperative dimerization, but the specificity towards AuxREs appears to be solely determined by the DBD (10, 12). Hence, the ARF DBD defines the auxin-responsive transcriptome.

The DBD structure can further be subdivided into a Dimerization Domain (DD), a B3 domain, and an Ancillary Domain (AD)(Fig. 1A)(10). The DD and AD fold together and tether the B3 domain. B3 domains form direct bonds with the AuxRE in the DNA, generally composed of TGTCNN elements (10, 13-16). Of these nucleotides, the TGTC forms the conserved core, while N5 and N6 are more variable. Among the AuxRE’s, TGTCGG elements show the highest ARF DNA-binding affinity. *Arabidopsis thaliana* (Arabidopsis) ARF1 (AtARF1) structure bound to an AuxRE indicated that four B3 domain residues are critical for AuxRE binding: R181, P184 and R186 bind to the TGTC core, and H136 binds to N5 and N6. Conformational flexibility of H136 and its flanking residues are thus responsible for the different affinities to TGTCNN elements (10, 14). The DD-AD scaffold enables ARF homodimerization in a head-to-head manner, allowing ARFs to bind bipartite AuxRE elements in an inverted repeat (IR) configuration. ARFs have also been shown to bind everted repeat (ER) and direct repeat (DR) bipartite motifs (17-19), likely requiring ARF homodimerization through a yet unknown mode of dimerization. Both AuxRE binding and bipartite element spacing preferences are thus completely determined by the DBD, although caused by different DBD subdomains.

**Figure 1.**
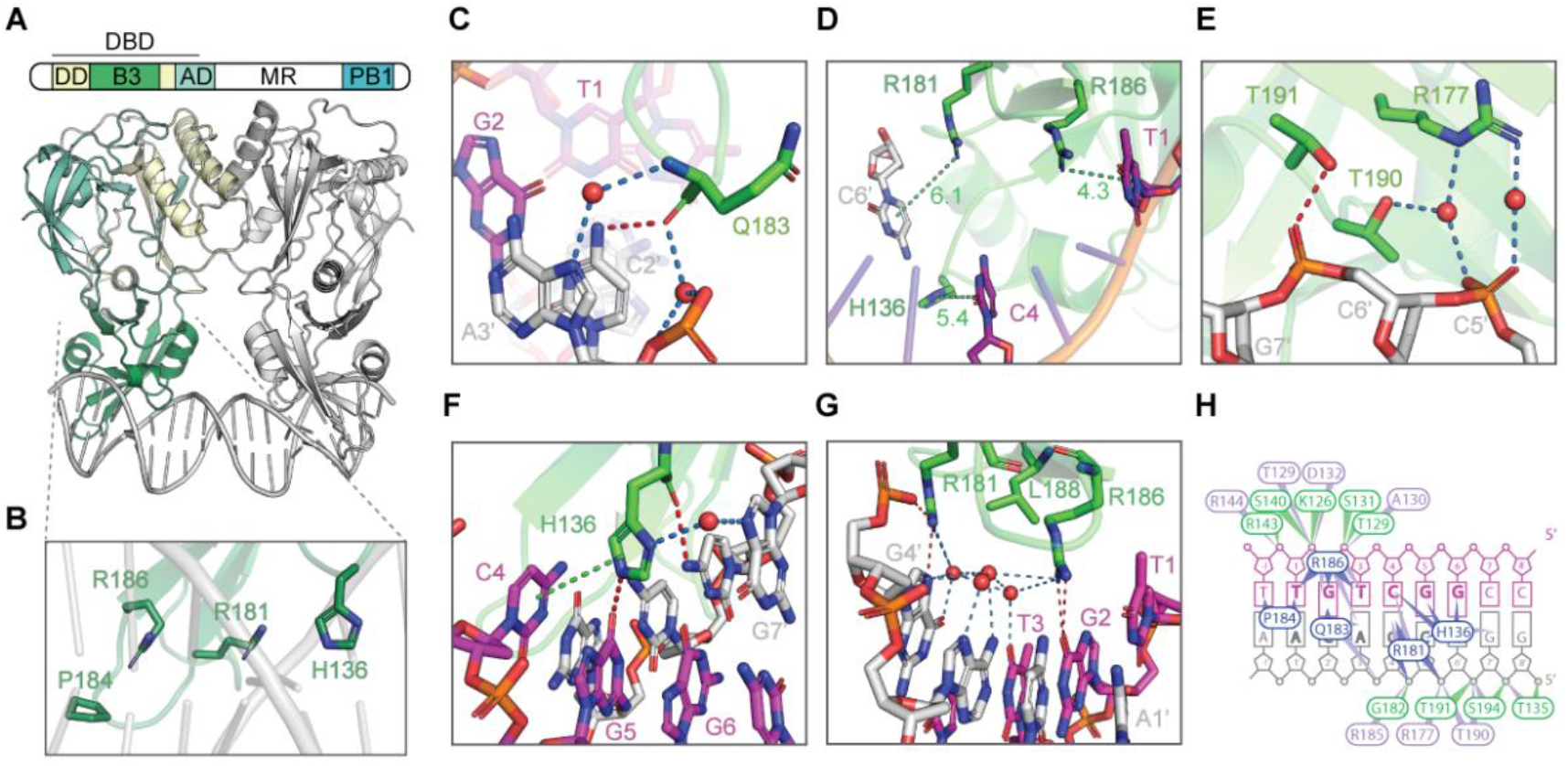
Extended ARF DNA-binding interface of AtARF1. **(A)** Anatomy of an ARF. Top panel shows the topology of a general ARF, the lower panel shows a cartoon representation of the AtARF1 DNA-binding domain (DBD) in complex with an AuxRE (PDB: 6YCQ). Dimerization Domain (DD; yellow), B3 domain (green), Ancillary Domain (AD; cyan). **(B)** Closeup of the B3-AuxRE interface represented as sticks, the main residues responsible for AuxRE specificity as identified by Boer et. al. (2014). **(C-G)** Closeups of the extended ARF- AuxRE interface of AtARF1. (C) Q183; (D) π-interactions of H136, R181, and R186; (E) water-mediated hydrogen bonds by R177 and T190; (F) π-interaction and water-mediated hydrogen bonds by H136; (G) water network directed by R181, R186, and L188. Interacting residues and nucleotides represented as sticks, with carbon molecules colored according to AtARF1 residues (green), DNA leading strand (magenta) and lagging strand (silver). Water molecules represented as red spheres. Non-covalent interactions (dashed lines) indicate π- interactions (green, with distance in Å in D), and hydrogen bonds between AtARF1 and AuxRE directly (red) or water-mediated (marine). **(H)** Schematic overview of AtARF1 residues (circles) and their interactions. Residues bind nucleobases in the major groove (blue), the phosphate backbone (green) or only make water-mediated contacts (purple).

The origin of ARFs can be traced back to a streptophyte common ancestor, and proteins sister to all ARFs can still be found in some extant streptophyte algae (20-22). During early streptophyte evolution, ARFs duplicated to form two separate clades, the AB- and C-ARF. Both clades are present in extant streptophyte algae such as the Klebsormidiophyceae and Zygnematophyceae. A second major duplication event involving the AB-class gene occurred later in a last common ancestor of land plants, forming the A- and B-class ARFs found in extant land plants. While ARFs have the same domain architecture, functional properties differ between classes (10, 18, 21, 23, 24). One such difference is found in the DNA-binding preferences: while A- and B-class ARF AuxRE-binding are highly comparable, their bipartite spacing preferences are only partially similar. Studies in the liverwort *Marchantia polymorpha* indicate that DBDs of both classes are interchangeable *in vivo* to some extent, indicating that these classes have partly similar DNA-binding preferences (22, 24). Contrarily, while C-class ARFs can bind similar TGTCNN elements as A- and B-class ARFs, they bind an expanded repertoire of bipartite spaced elements *in vitro* and correspondingly, the *M. polymorpha* C-class DBD is functionally different *in vivo* (24, 25). This suggests that C-class DNA binding is highly divergent from the A- and B-classes, in line with studies indicating C-class ARFs perform functions unrelated to auxin signalling.

The ARF family has deep roots, and ARFs are essential to auxin-dependent development across land plants. Structures of Marchantia and Arabidopsis ARF suggest a conserved mechanism of dimerization and DNA binding. However, a key unanswered question is to what degree the protein-DNA interface has evolved to accommodate new function for auxin in more complex response systems. Here, we re-evaluate the ARF-AuxRE interface and identify all DNA-contacting residues. We show that variation for the DNA-binding residues within this interface is rare across the plant kingdom. We find that any variation in this interface impairs function in a model A-class ARF *in vivo*. We finally infer that the ARF B3 domains are the sole drivers of AuxRE-binding specificity and show a deep class-specific functional conservation.

## Results

### Identification of the ARF DNA-binding interface

Previous crystallographic studies identified four AtARF1 residues that enter the major groove and are responsible for ARF DNA-binding specificity – AtARF1-H136, R181, P184 and R186 (Fig. 1B)(10, 14). These residues were conserved between AtARF1 and MpARF2 structures (24). Because the most recent AtARF1 structure (PDB: 6YCQ) by Freire-Rios *et. al*. (2020) was of such resolution that it allowed to resolve individual atoms, including water molecules, we revisited the ARF-AuxRE interface in search for additional features of DNA-binding. Our analysis identified the carboxyl group in Q183 to form hydrogen bonds with nucleobase C2’ (Fig. 1C). This interaction is conserved in the homologous residue of MpARF2 (Q225; Fig. S1A). We also found that H136 forms π-π interactions with nucleobase T1, while R181 and R186 form π-cation interactions with nucleobases C4 and C6’, respectively, in the AtARF1 structure (Fig. 1D). In MpARF2 however, only R225 (homologous to AtARF1-R186) makes pi-cation interactions (Fig. S1B). Pi-interactions therefore likely contribute to ARF AuxRE-binding.

In addition to direct interactions between ARF-residues and DNA-nucleotides, water molecules can be used as ‘bridges’ for residues to form indirect hydrogen bonds with the DNA (26). These water-mediated hydrogen bonds were also present in the AtARF1 structure. We observed, for example, two water molecules in the crease between T190 and R177 and the phosphate backbone that act as ‘bridges’ (Fig. 1E). A total of eight residues forming water-mediated hydrogen bonds are present at the periphery of the ARF-AuxRE interface: T129, A130, D132, R144, R177, R185, T190 and S194. Furthermore, base recognition by a TF can be achieved without direct contacts between residues and nucleobases by water molecule-networks (27, 28). In line, we observed water molecules in the space between the ARF and the major groove that enable water-mediated hydrogen bonds. Q183 forms water-mediated hydrogen bonds to the phosphates of C2’ and nucleobase A3’ (Fig. 1C). H136 forms these bonds with G7’, a nucleobase outside the TGTCGG element (Fig 1F). In comparison, R181 and R186, form a more extensive network of four water molecules, creating water-mediated contacts with T3, A3’ and G4’. A hydrophobic residue, L188, pointed towards the DNA and possibly repels the hydrophilic groups of the water molecules and the flanking R181 and R186 residues to increase the stability of the network (Fig. 1G). Overall, our reanalysis demonstrates the existence of previously overlooked set of atomic interactions in an extended ARF-AuxRE interface (Fig. 1H).

### Ultraconservation of the DNA-binding residues and B3 domains of ARFs

The extensive interactions between ARFs and AuxREs, and the finding that ARFs from all classes and different species (e.g., maize, Arabidopsis, *M. polymorpha, Chlorokybus melkonianii*) bind TGTCNN elements (8, 10, 18, 21, 24), suggest that the DNA-binding residues should be strongly conserved. We analysed the conservation of individual residues in the AtARF1 structure using ConSurf (29). The mean conservation score of the entire DBD is slightly above average conservation levels (5.47 ± 2.92, mean ± SD) (Fig. 2A). However, the B3 domain residues showed a much stronger conservation (7.52 ± 1.95), especially when compared to the rest of the DBD such as the DD-AD domains (4.65±2.84)(Fig. 2B). In fact, the DNA-binding residues were amongst the highest conservation scores (Table S1).

**Figure 2.**
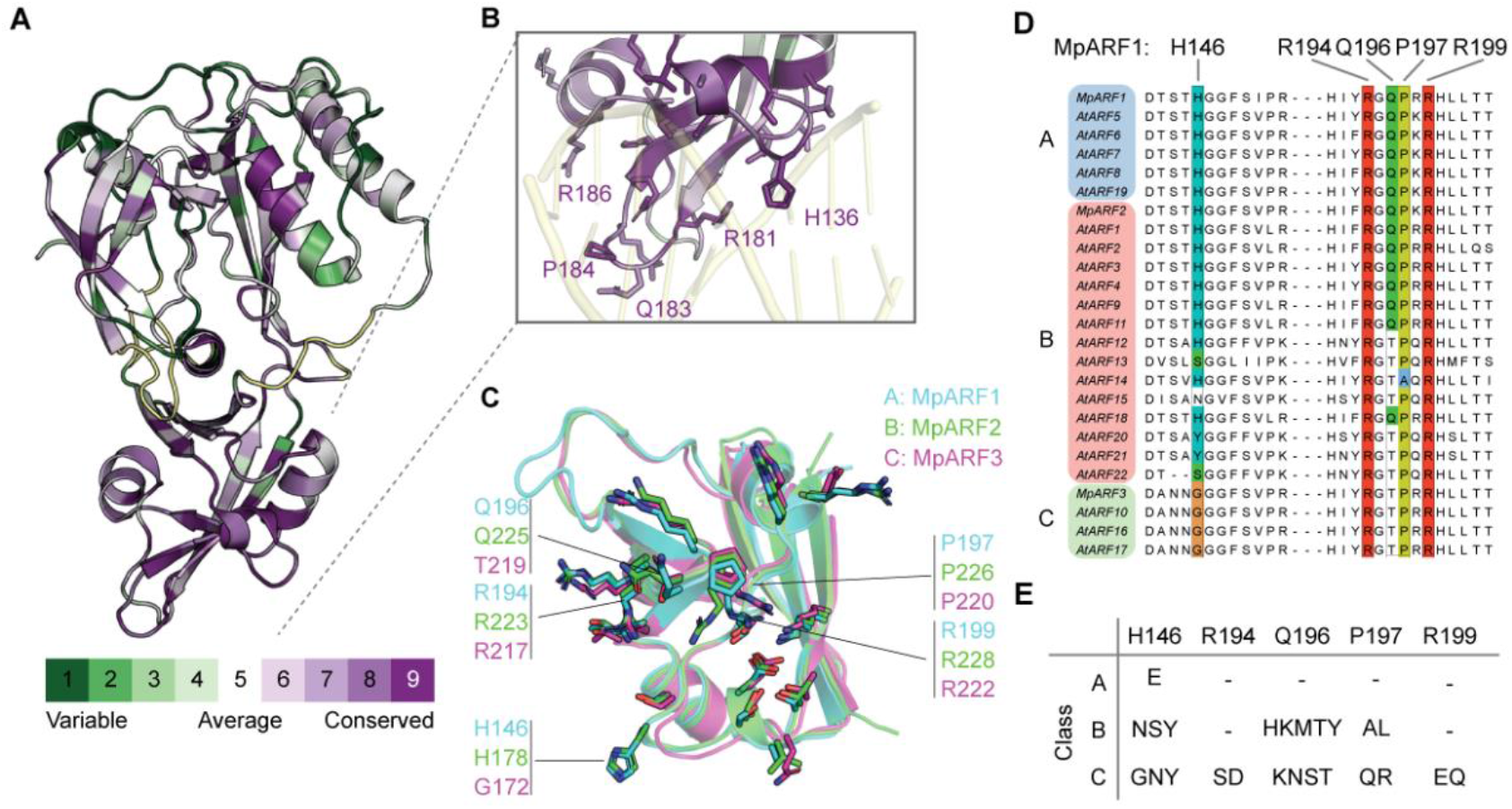
Conservation of ARF residues and structure. (A & B) Conservation analysis of AtARF1 residues using ConSurf (Ashkenazy et. al. 2016). Individual residues are colored according to their conservation score ranging from variable to conserved (1-9). (A) Whole DBD. (B) Closeup of B3 domain. DNA-interacting residues from Fig. 1 are shown as sticks. **(C)** Alignment of predicted protein structures of M. polymorpha ARF1 (MpARF1, cyan, A-class), MpARF2 (green, B-class) and MpARF3 (magenta, C-class) using AlphaFold2 (30). The AtARF1 homologous DNA-binding residues are represented as sticks, with major groove-binding residues in each ARF indicated. **(D)** Alignment of the DNA-binding loops from M. polymorpha and Arabidopsis ARFs. Colored are the major groove binding residues of MpARF1. **(E)** Table with all variation found in the land plant A, B and C-class ARFs from the phylogenetic tree of Mutte et. al. (2018).

A prerequisite to achieve proper DNA-binding would be that the key residues acquired the same 3D location in the structure. We investigated this by predicting the protein structures of the three *M. polymorpha* ARFs, each belonging to one of the major land plant classes, MpARF1, MpARF2, and MpARF3 (A, B, and C, respectively)(Fig. 2C), using AlphaFold2 (30, 31). All key residues showed an identical position in the predicted structure, corroborating previous homology models (20, 21, 24). This indicates that all ARFs have a highly similarly folding B3 domain, with key DNA-binding residues in the same 3D location.

The conservation of the DNA-binding residues and DNA-interaction interface suggests that there is a strong evolutionary pressure to preserve ARF DNA-binding specificity. We wondered if any functional variation is found in these residues, for example as part of diversification of auxin-dependent gene regulation in land plant evolution. We first examined variation within a collection of naturally occurring *M. polymorpha* accessions (32). Each ARF class gene has a single ortholog in all these accessions, and thus, variation should be restricted or absent if ARF specificity is selected for. Indeed, the B3 domain protein sequences of MpARF1 and MpARF3 are entirely conserved in all 133 sequenced accessions. For MpARF2, several accessions have a MpARF2-N203S polymorphism, a residue not involved in direct DNA-binding. Therefore, *ARF* genes do not tolerate variation and experience selection pressure to maintain the protein sequence in *M. polymorpha*. In contrast, Arabidopsis contains several paralogues genes for each ARF class (i.e.: 5, 15 and 3 copies of A, B and C-class respectively), with overlapping, redundant and opposite functions (22, 24, 33-36). Redundancy in paralogous functionality in TF families could allow for mutations that might lead to sub- or neofunctionalization of DNA-binding (37, 38). We therefore assessed whether mutations occurred in DNA-binding residues in ARF genes in Arabidopsis. Beyond the Histidine-Glycine variation (20, 21), variation does not occur within A and C-classes, suggesting divergence between paralogs does not involve DNA-binding specificity changes (Fig. 2C). The main B-class paralogs, like AtARF1, similarly retain these residues. However, the recently expanded subclade of Brassicaceae-specific B-class ARFs on chromosome 1 (ARF12-15, 20-22)(36, 39) show variation in three of these residues, with AtARF1-Q183 being substituted by Thr in all instances, H136 showing substitution for either Asn, Ser or Tyr, and P184 being substituted for Ala only in AtARF14. To further explore the idea of genetic variation being almost absent, and permitted only in redundant paralogs, we re-analysed a list of previously collected, curated and phylogenetically positioned ARFs (Mutte et al. 2018). This list contains 258, 280, and 123 ARFs of the A, B, and C-classes, respectively, covering all clades off land plants. We then analysed the variation existing among the DNA-binding residues (Fig. 2E, Table S1). Among major groove binding residues, we found a single Glu variant instance at MpARF1-H146 position in one of the three AtARF6 homologues of the magnoliid *Calycanthys floridus*, indicating less than 0.4% variation of this residue among A-class ARFs. Among B-class, H146 showed the highest variation, caused by the above-mentioned diverging Brassicaceae-subclade, apart from a single instance of variation in this residue outside this subclade (Glu, *Hakea prostrata*, Proteaceae).

Q196 was the most variable position in the B-class ARFs, with 19% (53) of the B-class ARFs having a histidine at this position. All these B-class ARFs are found in mosses, lycophytes and ferns and are not limited to a single B-class clade, indicating independent mutations are causal for this variant. The physiological implication of this mutation is unclear, especially since Q196 is somewhat tolerant to mutations given its mode of base interaction, yet no variation is found in this residue in other ARF classes even when redundant paralogs exist.

Finally, the glycine at position H146 is conserved in virtually all C-class ARFs. All the variation summarized in Fig. 2E for the C-class ARFs were derived from four C-class ARFs: rice OsARF20, a putative pseudogene with two DBDs (40); and maize ZmARF31, 32 and 33.

These results highlight that ARF DNA-binding residues are highly intolerant to variation, and this variation only occurs in diverging paralogs possibly undergoing sub-/neo-functionalization of the DNA-binding mechanisms, or under pseudogenization mechanisms. Altogether, this indicates that most variation in DNA-binding residues likely leads to reduced protein function, and provides a hypothesis to why these residues are ultra-conserved.

### *M. polymorpha* sole A-class ARF function depends on the core TGTC-binding residues

To analyse the physiological relevance of the AuxRE binding residues, and to bypass the inherent restrictions of using genetic systems with multiple redundant paralogous ARFs, we focused on *M. polymorpha* single A-class ARF. MpARF1 has been proposed as the only positive effector of auxin-induced transcription, and the Mp*arf1* mutants do not respond to exogenous auxin (Fig. 3A) (41, 42). The auxin insensitivity of the null mutants can be complemented by the introduction of the native coding sequence (24, 42). However, it is unclear if there are additional auxin-dependent effectors mediating auxin transcriptional responses. To explore this possibility, we performed whole-transcriptome sequencing analyses of wild-type (Tak-1) and MpARF1 null mutant (Mp*arf1-4*) in response to a short-term treatment with the auxin indole-3-acetic acid (IAA) after a prior depletion of the endogenous auxin through treatment with biosynthesis inhibitors L-Kynurenin (L-Kyn) and Yucasin. Tak-1 responded to IAA with 112 genes being differentially expressed (69 up, 43 downregulated). In contrast, the Mp*arf1* mutant showed no response to auxin (Fig. 3B). This indicates that MpARF1 is the only effector mediating auxin-induced transcription in *M. polymorpha*.

**Figure 3.**
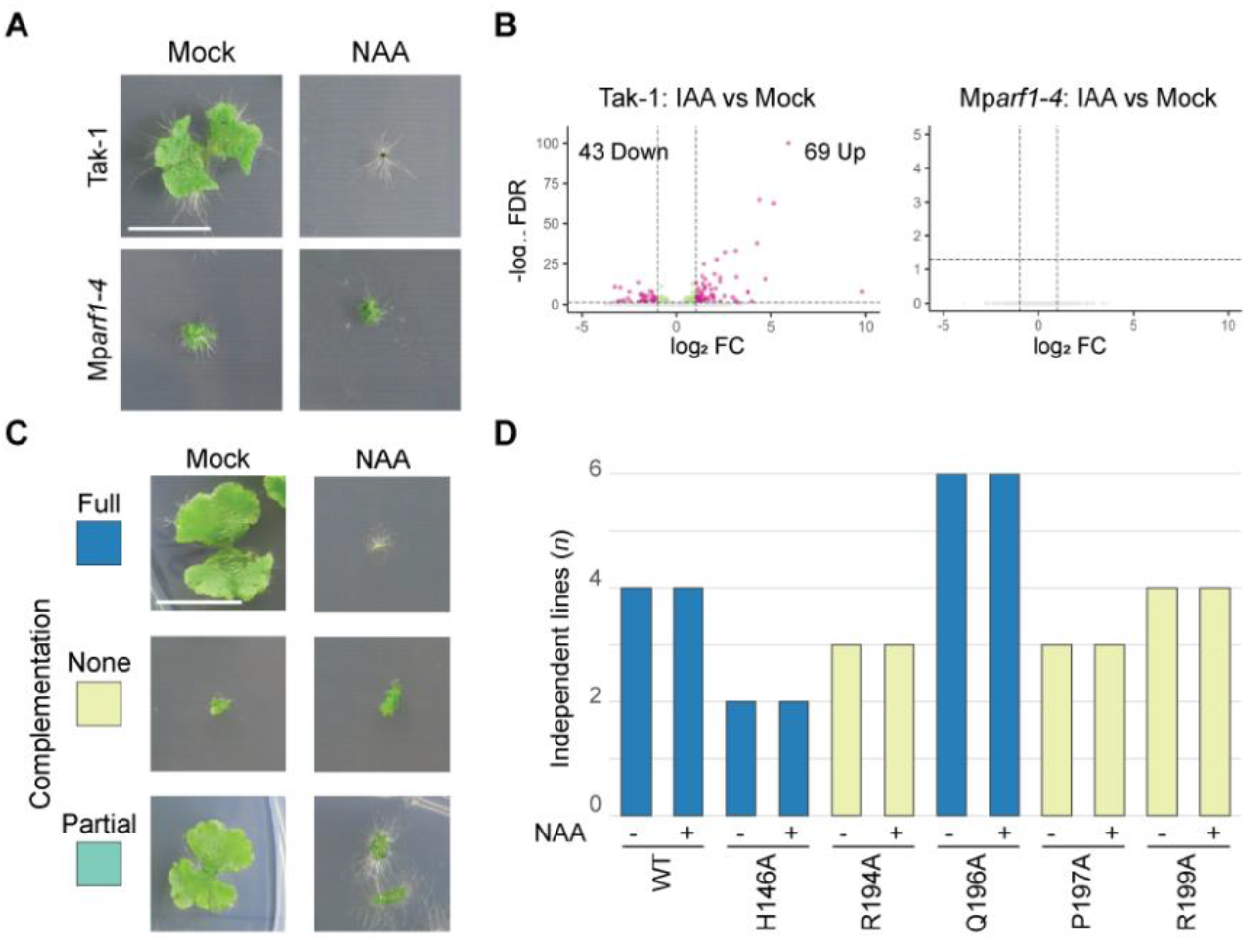
Conserved DNA-binding residues are critical for the function of MpARF1 in vivo. **(A)** 14-day-old plants of Tak-1 and Mp*arf1-4*, grown on mock (DMSO) or auxin (3 μM NAA) medium. **(B)** RNA-seq volcano plots of differentially expressed genes (DEGs) in Tak-1 and Mp*arf1-4* gemmae L-Kyn and Yuc treatment, followed by auxin (IAA). False Discovery Rate (FDR); Fold Change (FC). **(C)** 14-day-old plants corresponding to full (blue), none (yellow), or partial (turquoise) complementation of the Mp*arf1-4* phenotypes on mock and auxin containing media. (A&C) Scale bar is 1 cm. **(D)** Schematic overview of phenotypes of 14-day-old plants of Mp*arf1-4* complemented with alanine substitution variants of MpARF1. Summary of experiments shown in Fig. S2 and S4A.

MpARF1 was previously shown to bind TGTCGG elements *in vitro* (24). In line, it shows full conservation of the key DNA-binding residues (Fig. 2C). We thus tested if these residues are required for the *in vivo* function of MpARF1 by performing alanine substitutions of H146, R194, Q196, P197 and R199. We used a complementation assay of Mp*arf1-4* to test whether the alanine variants were able to fully complement, partially complement or could not complement growth impairment under standard conditions or the auxin response when grown on medium with the synthetic auxin 1-Naphthaleneacetic acid (NAA; Fig. 3C). The alanine versions R194A, P197A and R199A were unable to complement the Mp*arf1* phenotypes, indicating these residues are essential for MpARF1 physiological function (Fig. 3D, Fig. S2, S4). The full complementation shown by H146A and Q196A mutants suggest these residues are not essential for function. Our structural analysis suggests that Q196 binds C2’ via its carboxyl group, and an alanine substitution would likely be able to retain this interaction and would therefore likely tolerate different amino acids at this position. In contrast, H146 was suggested to be critical for binding high affinity TGTCGG elements, yet the H146A mutant exhibits full complementation. This data highlights the requirement of R194, P197 and R199 in conferring ARF functionality, and the deeply conserved role of their base-specific binding to AuxRE that endow ARF-dependent auxin responses.

### Natural variation in TGTC-binding residues impairs the function of MpARF1

Given the deep conservation found in the ARF DNA-binding in plants, the limited variation in any of the key DNA-contacting residues is likely to affect DNA-binding properties. This could either generate specific ARF functions, unique to the context in which the variant is found, or may reflect loss of function mutations. The latter is not unrealistic, given that most variation is found in paralogous duplicates. To address function of the ARF variants, we assessed the ability of a series of mutations to complement Mp*arf1* mutant phenotype. We found R194 and R199 to have extensive interactions with the AuxRE (Fig. 1G) and therefore reasoned that the observed variation in the C-class ARFs (SDEQ) would likely disrupt AuxRE-binding (Fig. 2E). Indeed, the charge reversal R199E mutant is unable to complement Mp*arf1* phenotypes (Fig_4A, Fig. S3). We therefore tested if the similar physiochemical lysine could substitute for R194 and R199, but neither R194K nor R199K were able to complement, making it unlikely for any other amino acid to substitute the function of both arginine’s in AuxRE-binding. For the Q196 residue, we made Q196K and Q196R substitutions that were both able to complement, showing that this residue could tolerate other long, polar amino acids. Finally, we tested P197Q and P197R substitutions based on the C-class ARFs, both of which were unable to complement, suggesting the position cannot tolerate polar residues. In addition to DNA-binding by P197 in the ARFs (10).

**Figure 4.**
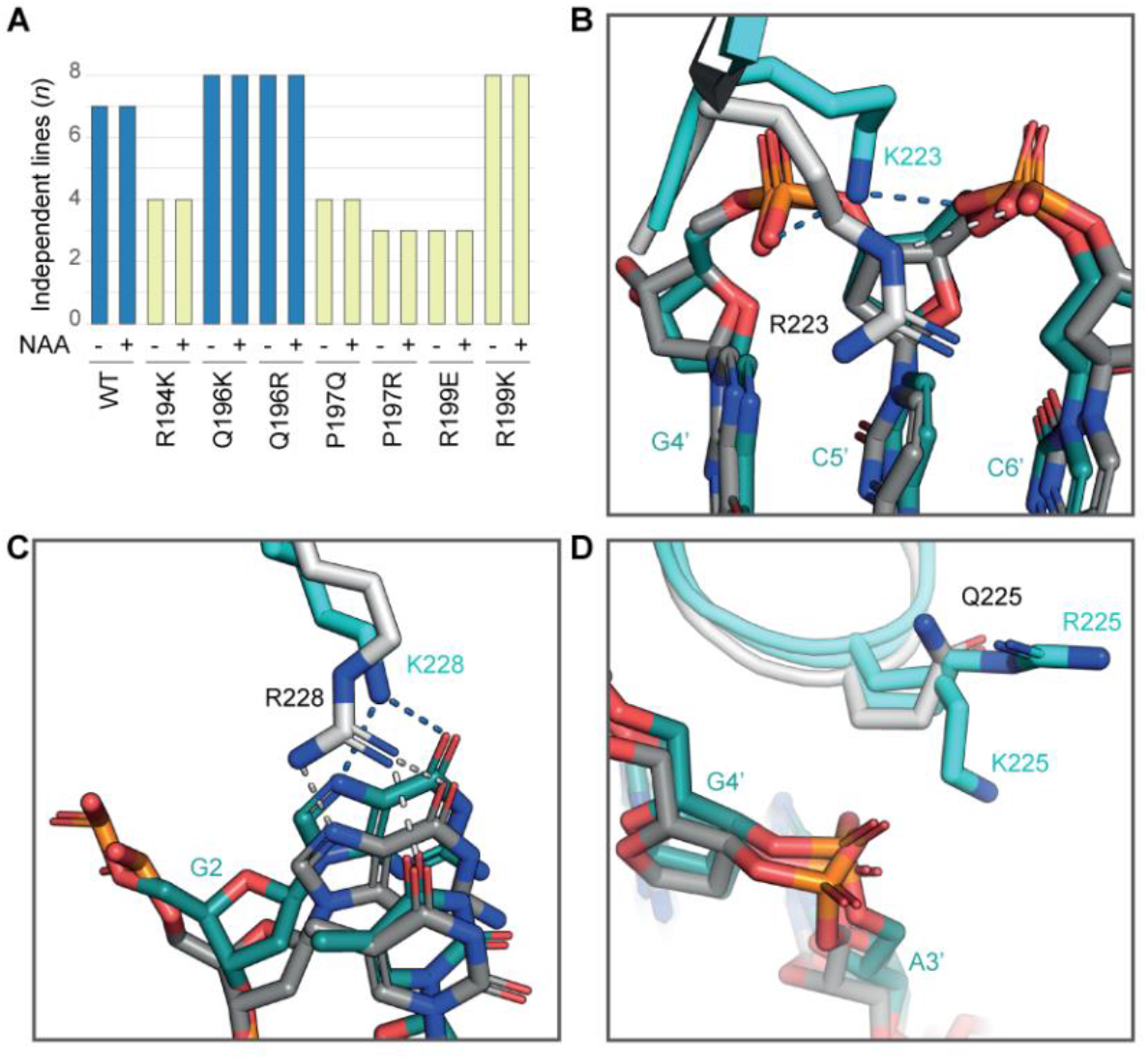
Variation for TGTC-binding residues impair MpARF1 function in vivo. **(A)** Schematic overview of complementation of 14-day-old plants grown on mock (DMSO) or auxin (3 μM NAA), based on experiments shown in Supp_CorevarPheno1. **(B-D)** Structural alignments of MpARF2 (PDB: 6SDG, white/grey) and MpARF2 variants (cyan/teal) R223K (B), Q225K/R (C), and R228K (D) after molecular docking on the original DNA. White/Blue dashed lines: hydrogen bonds in original and docked structures respectively. MpARF2-R223, Q225, and R228 correspond with MpARF1-R194, Q196, and R199 respectively.

To understand how these residue variants may influence ARF DNA-binding, we performed *in silico* molecular docking. We used the solved MpARF2 crystal structure (PDB: 6SDG), changed the corresponding residue, and docked the ARF variant with TGTCGG DNA. While the native R223 (MpARF1-R194) enters deeply into the major groove, the docked K223 variant could only reach the phosphate backbone (Fig. 4B). Similarly, whereas R228 (MpARF1-R199) hydrogen bonds with G2 and T3, K228 only bonded with G2 (Fig. 4C). R194 and R199 are therefore crucial residues for ARF DNA-binding. In contrast, the docked K225 and R225 variants of Q225 (MpARF1-Q183) do not form additional contacts and point outwards of the major groove.

Together, these results confirm the crucial role of R194, P197 and R199 for ARF DNA-binding and suggest that their physiochemical role in DNA-binding cannot be replaced by other amino acids. On the other hand, while Q196 is almost fully conserved in the A-class ARFs, this residue tolerates alternative chemical and structural amino acids, suggesting it may fulfil an additional essential role, possibly unrelated to DNA-binding. Hence, we conclude that the observed variation in ARF DNA-contacting residues is non-functional and that the protein-DNA interface is under extremely strong purifying selection.

### A single, tunable residue in the ARF-DNA interface

While variation in DNA-contacting residues appears non-functional, there is one interesting exception. The Histidine that is conserved in A-class and B-class ARFs is replaced by a Glycine in C-class ARFs. The MpARF1-H146 residue does not bind the core TGTC-site but is thought to determine the affinity towards N5 and N6 of TGTCNN elements (10, 14). We identified several alternative residues at this site in the land plant ARF family (Fig. 2E), and it is possible that variation at this one site modulates ARF DNA binding specificity and biological function. We used our genetic model to study the functional relevance of the full set of variants found in nature (H146G/N/S/Y) (Fig. 5A). We also included a set of mutations to test the relevance of this residue for DNA binding and explore the range of physicochemical properties that are compatible with its function. To this end, we engineered a H146A variant as part of an Alanine scanning. We also engineered a H146F as an aromatic, non-polar residue with a size similar to Histidine, and H146Q as a polar, non-charged residue. Lastly, we engineered a charge reversal mutation (H146E) that would be predicted to strongly disturb the mechanism of DNA interaction. MpARF1 carrying the H146E mutation was unable to complement, confirming the critical relevance of this residue for DNA binding (Fig. 5A, S4, S5). Interestingly, all other variants were able to fully or partially complement a range of *arf1* mutant phenotypes. All variants except H146E were able to at least partially restore growth after 14 days on medium with or without auxin (Fig. 5A-B; Fig. S4,S5), while thallus area after 35 days of growth were comparable to Tak-1 for all H146 variants (Fig. 5C-D). Additionally, all the tested variants except the H146G variant produced similar amounts of gemma cups compared to Tak-1 (Fig. 5E). Finally, while Mp*arf1-4* does not produce antheridiophores after 5 weeks of far-red light treatment (Fig. 5F), all H146 variants restored antheridiophore production. These results suggest that the H146 residue could allow for evolutionary tuning.

**Figure 5.**
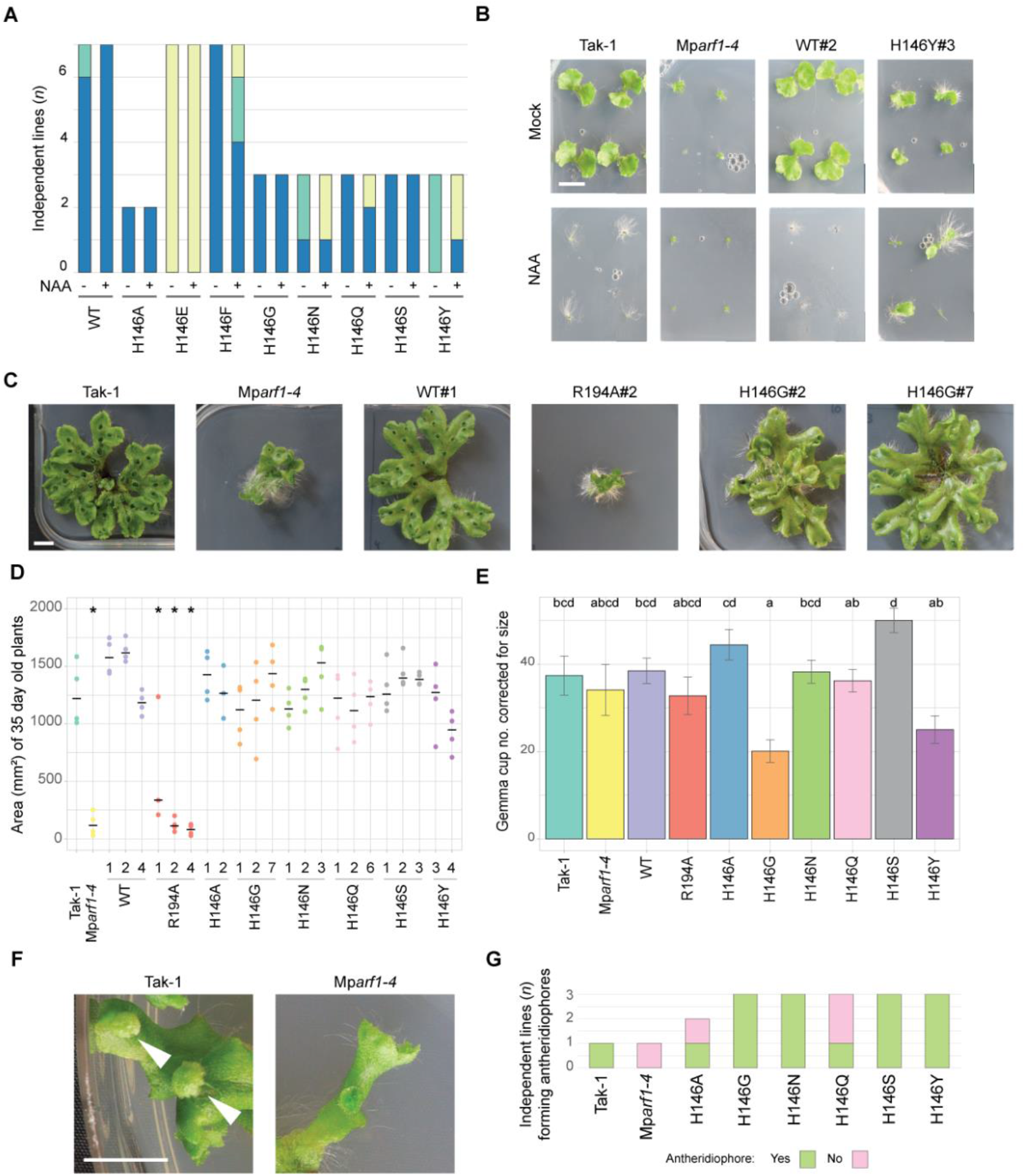
Variation for TGTC-binding residues impair MpARF1 function in vivo. **(A)** Schematic overview of the number of independent lines that complemented the growth phenotype of 14day-old plants. Based on phenotyping data in Fig. S4. **(B)** Pictures of 14 day-old plants of Tak-1, Mp*arf1-4*, WT#2 and H146Y#3 corresponding to full and partial complementation by the WT and H146Y constructs respectively. See also Fig. S4. **(C)** 35 day-old plants. **(D)** Area (mm^2^) of 35 day-old plants (*n* per line = 4). Horizontal lines are mean per line. Asterisks indicate lines with statistical significant differences to Tak-1, corresponding to p ≤ 0.05 following ANOVA and posthoc testing (Sidak correction). **(E)** Gemma cup per genotype, corrected by thallus size (ANCOVA). Error bars are SD. Small letters correspond to different statistical groups following posthoc testing (SIDAK correction, p ≤ 0.05). **(F)** Closeup of five week-old Tak-1 and *Mparf1-4* thallus. Arrows indicate antheridiophores. **(G)** Number of independent lines per genotype that formed antheridiophores after five weeks.

These results suggest that MpARF1-H146 has a unique character in the ARF family: it is critical for biological function, yet allows limited variation in the amino acids that can occupy this position. We explored this further by analysing in detail the single-most prevalent residue at this position. All C-class ARFs have a Glycine at this position, but the impact of this variation has not been described. The H146G variant was also able to complement MpARF1 function in thallus growth, antheridium development and auxin response (Fig. 5A, 5D, 5G). However, we noticed that plants expressing the H146G variant developed fewer gemmae cups (Fig. 5C, 5E). This clearly demonstrates a functional difference in biological function between the Histidine and Glycine residues in the ARF-DNA interface.

### B3 domain functions are conserved across land plants

The deep conservation of the key ARF DNA-binding residues, and their fundamental role in ARF physiological function suggests that the DNA-binding interface represents an ancient feature under strong purifying selection. If true, one would expect that the homologous DNA-binding interface among distantly related ARFs should be functionally interchangeable. We tested this hypothesis by replacing the entire B3 domain in MpARF1 by orthologous domains form each Arabidopsis A-class ARF. These 5 AtARFs (AtARF5,6,7,8,19) represent duplicates that emerged in the common ancestor of vascular plants, seed plants and flowering plants, and may represent subfunctionalized, redundant or neofunctionalized copies. All these A-class ARFs shared the same DNA-binding residues (Fig. 6A) that adopted the same 3D orientation in homology models (Fig. S6). All the AtARF swapped versions completely complemented the Mp*arf1* phenotype, indicating they all share the same binding preferences (Fig. 6B, S7). This finding suggests that no functionally relevant divergence has occurred in the B3 domain of these ARFs since their split from the land plant common ancestor.

**Figure 6.**
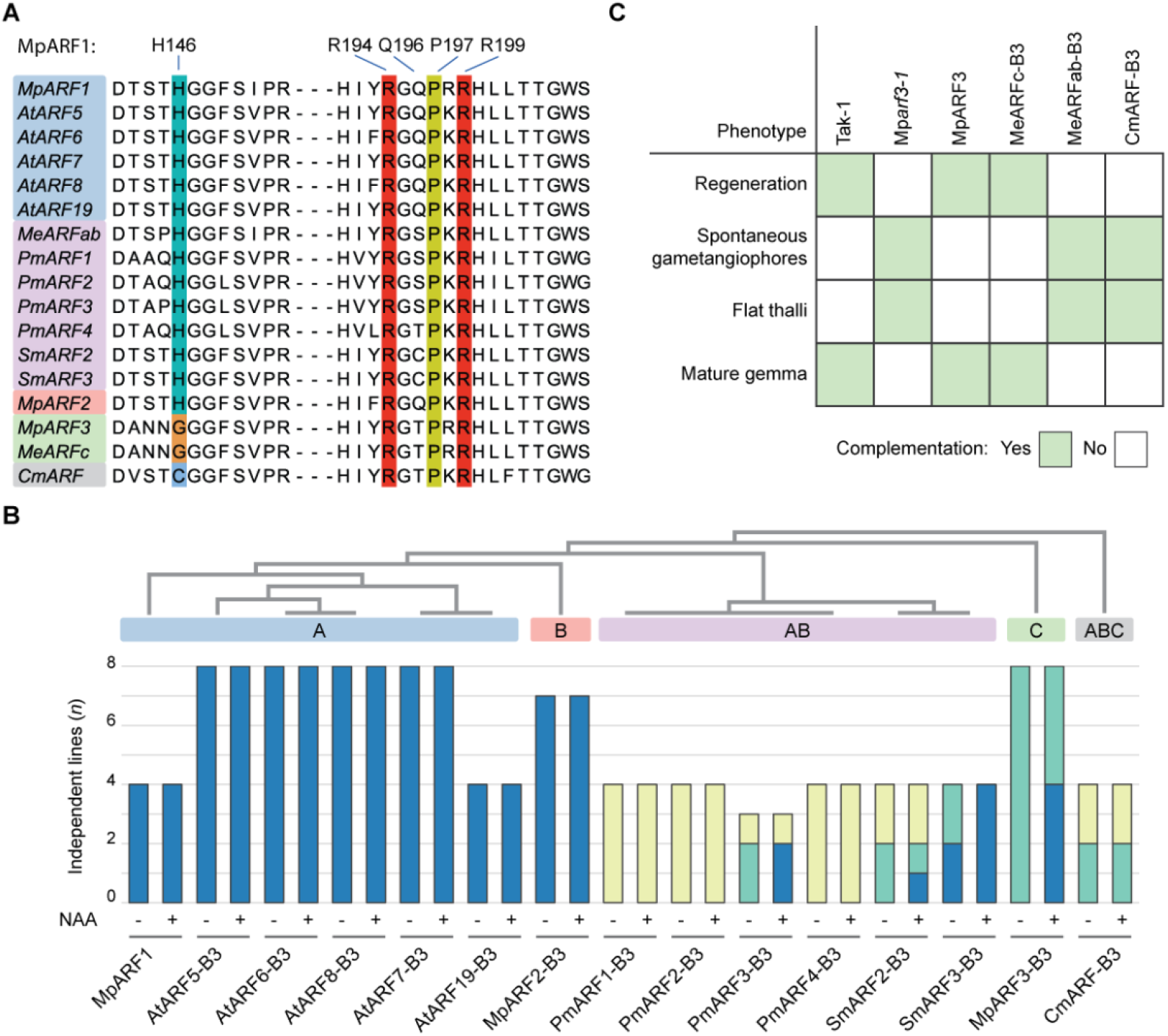
B3 swaps with different B3 domains. **(A)** Alignment of ARFs used for B3 swap experiments. **(B)** Schematic overview of number per lines for each B3 swap that complemented Mp*arf1* phenotypes. Top, schematic phylogenetic tree showing relationship of each ARF and each ARF-class. Lines were grown for 14 days on medium without (-) or with (+) NAA. All lines represent Mp*arf1-4* transformed with *p*Mp*ARF1::*Mp*ARF1* with the endogenous B3 (MpARF1) or with B3 domain swapped (rest). **(C)** Schematic overview of complementation of Mp*arf3* phenotypes by B3 swap constructs.

Auxin response in *M. polymorpha* hinges on competition in DNA binding between the A- and B-class ARF (24). This is based on common DNA-binding specificity of A- and B-ARFs, likely inherited from an ancestral AB-type ARF (22). However, replacing the MpARF1 DBD with that from MpARF2 does not fully restore function (24). Since the DBD acts both as DNA-binding unit and as dimerization unit, we explored the functional equivalence of only the DNA-binding unit: the B3 domain. The MpARF2-B3 swap was fully able to complement the Mp*arf1* mutant phenotype (Fig. 6B, S8), confirming identical intrinsic DNA-binding properties. As a control, we swapped the MpARF3 B3 domain into MpARF1, and found poor complementation (Fig. 6B, S8). Thus, MpARF1 and MpARF2 share equivalent B3 domains, while the one from MpARF3 is functionally different.

We can thus infer that the MpARF1/2 binding specificity emerged at least as early as in the A/B ancestor. However, we recently reconstructed the evolutionary history of the ARF proteins (22), and identified extant copies that may reflect earlier ancestral states. We therefore swapped the MpARF1 B3 domain with a series of extant algal B3 domains that represent AB or ABC groups. We decided to use the streptophytes *Penium margaritaceum, Spirogloea muscicola* and *Chlorokybus melkonianii*. Penium and Spirogloea are part of the closest sister lineage to land plants, the Zygnematophyceae, while Chlorokybus represents an earlier split from the lineage that generated land plants. *S. muscicola* has a single AB-ARF ancestral gene copy that has triplicated into three similar paralogues. In contrast, *P. margaritaceum* is part of the Desmidiales, a derived group within the class, and its four AB-ARFs represent two rounds of duplications occurring hundreds of million years ago, thus allowing for sub- and neo-functionalization events. All these AB-B3 domains have conserved DNA-binding residues, except of the residues at position Q196 (Fig. 6A). While PmARF3-B3 was able to partially complement the Mp*arf1* phenotypes, the remaining PmARF-B3’s were not (Fig. 6B, S9, S10), despite conservation of the critical DNA-binding residues (Fig. 6A). SmARF2-B3 was similarly poorly complementing. In contrast, all lines of the SmARF3-B3 swap had a fully complemented Mp*arf1* auxin phenotype, while two of four lines showed full complementation of growth under mock conditions, indicating that the ancestral AB-ARF must have had similar DNA-binding preferences as extant A/B-class ARFs from land plants. The single ARF of *Chlorokybus melkonianii* derives from an ancestral ABC-class ARF, predating the split of AB and C-classes (22). Strikingly, the CmARF-B3 was able to partially complement Mp*arf1*, indicating that the ABC-class DNA-binding is similar to that of A-class (Fig. 6B). This suggests that DNA-binding preferences are similar among the majority of ARF classes, the exception being C-class.

Our previous analyses indicate that C-class DBDs diverged significantly from AB-class DBDs, while ABC-class DBDs have a different functionality than both classes (22). To study if ABC- and C-class binding preferences are comparable, we performed complementation assays using B3 swaps with the Mp*arf3* mutant. None of the phenotypes associated with the mutant were complemented when using B3 swaps from non-C-class ARFs, including CmARF-B3 (Fig. 6C). In line with deep conservation of C-class, the *Mesotaenium endlicherianum* ARFc-B3 (MeARFc-B3) swap restored the phenotype on the mutant, unlike the MeARFab-B3. Overall, this suggest that the DNA-binding preferences greatly diverged within the C-class from the ancestral state, while it was partially retained in AB-class.

## Discussion

Auxin-dependent gene regulation is key to its roles in growth and development, and is mediated by DNA-binding ARF proteins. The protein-DNA interface is therefore the ultimate effector of specificity in auxin-dependent gene expression. A key question is if and how this interface has evolved. We address this question here and find the interface to be ultraconserved and intolerant to variation.

We first explored whether, apart from the singe A-class MpARF1 protein, other transcription factors may mediate auxin-dependent gene expression in Marchantia. Transcriptome analysis on an *Mparf1* mutant revealed that no differential gene expression could be detected in its absence, and this suggests that MpARF1 is the only transcription factor transducing the auxin signal. This is consistent with previous studies, where domain swapping of the PB1 domain of MpARF1 with MpARF2 was able to partially rescue the growth phenotype of Mp*arf1-4*, but not its response to exogenous auxin (24). We thus consider MpARF1 as a strong model system to study the core functions of A-class ARFs across land plants. We here used a series of computational and genetic assays to explore the structure-function relationship and evolvability of the ARF-DNA interface.

First, we replaced the entire B3 DNA-binding subdomain in ARF1 with the one from other ARFs. Earlier swaps of the entire DNA-binding domain between MpARF1 and MpARF2 indicated that MpARF2-DBD was not completely interchangeable with MpARF1’s (24). In contrast, we found that their B3 domains were fully interchangeable, suggesting that the differences between MpARF1 and MpARF2 DBDs are found in the DD-AD domains, probably attributable to spacing preferences or co-factor recruitment. The MpARF3 DBD failed to complement the Mp*arf1-4* mutant (22, 24), while here we found that the B3 domain can partially complement the mutant phenotypes although poorly. These results indicate that MpARF3 is different with respect to both DNA-binding specificity as well as spacing preferences. These new results strongly support the notion that A- and B-ARFs retained the DNA-binding specificity of the ancestral AB-ARF, and this underlies the competitive DNA binding by the two classes.

The ARF B3 domain is conserved across the entire streptophyte lineage. We showed here that mutations in its DNA-binding residues impair the biological function of MpARF1. If mutations are so detrimental for their function, it explains why variation in DNA-binding residues is found only in species with expanded families of ARFs. Especially mutations on the R194, P197 and R199 residues are detrimental. For other TF families, after gene duplication, subspecialisation can occur on DNA-binding preferences either by acquiring mutations that decrease binding affinity to the ancestral DNA-element, or increase binding to previously low affinity sites (43-45). For ARFs, *in vivo* function is immediately lost upon mutations of DNA-binding residues, suggesting that the family has a limited capacity to gain binding to novel elements, leading to rapid pseudogenization. This could explain why the ABC- and C-class ARF-B3s can still partially complement Mp*arf1* phenotypes and thus retain some DNA-binding. Whether the ARFs we observed with mutations on DNA-binding residues are truly becoming pseudogenes or whether they acquired a novel function in a context we did not test remains to be investigated in the corresponding species, or by B3 swap experiments as performed in this study. Together these results highlight the ultraconservation of the ARF DNA-binding preferences.

The B3 domain swap experiments here reveal how the ARF DNA-binding preferences evolved. The proto-AB-ARF likely acquired DNA-binding preferences that are still maintained in extant AB-class ARFs from streptophyte algae, as well as within the land plant A-class and B-class ARFs (22). Even in expanded A-class ARF families like Arabidopsis, we find that a conserved DNA-binding preference has been maintained. Considering the assumption that *M. polymorpha* likely reflects part of the ancestral function prior to subfunctionalization within the ARF family, we cannot exclude that the Arabidopsis A-class ARFs acquired additional DNA-binding in addition to their conserved DNA-binding. Furthermore, we only tested a single B-class and C-class ARF of *M. polymorpha* in this assay. Although DAP-seq results with maize ARFs (18) suggest that all B-ARFs share similar high-affinity DNA-binding preferences, whether they are all interchangeable could be achieved through similar experiments using an Mp*arf2* mutant. Finally, we only tested four algal AB-ARFs, of which two complemented, and two did not. The ones that did support the hypothesis that AB-ARFs have the same DNA-binding as A and B-class ARFs. The two that were unable could suggest sub- or neofunctionalization, or pseudogenization within the algal species.

In summary, our computational and genetic analysis of the ARF protein-DNA interface reveals an extreme degree of conservation that can likely explain the conserved roles of auxin siganling in growth and development of land plants.

## Material & Methods

### Plant materials and growth conditions

The male accession of *M. polymorpha*, Takaragaike-1 (Tak-1, male), the ARF null mutants Mp*arf1-4* and Mp*arf3-1* in Tak-1 background (20, 42) and generated lines were axenically and vegetatively propagated on half-strength Gamborg’s B5 medium (hereafter ½ B5; G0209, Duchefa Biochemie) pH 5.5-5.8 containing 1% agar (hereafter solid ½ B5) under 50-60 μmol photons m^-2^ s^-1^ constant white fluorescent light at 22°C.

### Plasmid construction

Oligonucleotides and plasmids used and generated in this study can be found in Table S2 and S3. Site-directed mutagenesis of H146, R194, Q196, P197 and R199 was performed by PCR-amplifying pENTr221_MpARF1 into linear fragments, followed by DpnI digestion (FD1703, Thermo Scientific), and by assembled using single strand Oligo (ssOligo) HiFi Assembly (E2621, New England Biolabs). B3 coding sequences were amplified from plasmids or cDNA and similarly assembled using HiFi Assembly or Golden Gate Cloning with BsaI (E1602, New England Biolabs). All entry vectors were subcloned into corresponding destination vectors via LR Clonase II (11791020, Invitrogen).

### Transformation of Mparf1-4 and phenotyping experiments

Mp*arf1-4* thallus material was transformed with binary vectors using an *Agrobacterium tumefaciens* (GV1301) protocol as described in Hernández-García *et. al*., 2024.

Growth experiments with or without NAA were performed by growing gemmae on solid ½ B5 with 3 μM NAA or without (adding an equivalent volume of DMSO) and growing at 50-60 μmol photons m^-2^ s^-1^ continues white fluorescent light at 22°C. After 14 days, pictures were taken with a Canon EOS250D camera with EF-S 18-55 IS STM lens. To quantify thallus growth area, pictures were resized to 2000×2000 pixels with ImageMagick (version 7.1.1-21), followed by pixel classification and object quantification of projected area of single gemmalings with Ilastik (version 1.3.3)(46).

For gemma cup number and 5 week growth experiment, gemmae were grown under the same conditions as above, except for 5 weeks. Projected thallus area was measured as above and gemma cup number counted manually.

To score antheridiophore production, a single gemmae per line was grown for 14 days at the conditions above, after which they were supplemented with far-red (FR) light (50-60 μmol photons m^-2^ s^-1^). Pictures were taken after five weeks at FR conditions.

### Transcriptome analysis

Plants used for RNA-seq were grown as follows: Gemmae of Tak-1 and Mp*arf1-4* were grown on solid ½ B5 as above. After 5-6 weeks of growth, gemmae were harvested from the plants by holding the plate upside down above a tray filled with sterile demineralized water and hitting the bottom of the plate with a metal spoon. Gemmae that fell in the water were collected in a 70 μm cell strainer. Gemmae were immediately submerged in a solution of liquid ½ B5 with 50 μM each of L-Kyn and Yuc. After 6 hours, gemmae were collected in a strainer and washed three times with sterile demineralized water. Gemmae were either submerged into liquid ½ B5 (mock) or liquid ½ B5 with 1 μM IAA (mock contained equivalent volume of DMSO). After 1 hour, gemmae were collected in 2 ml Eppendorf tubes, frozen in liquid nitrogen and ground into a powder with a bead shaker. RNA was extracted with the RNeasy Plant Mini Kit (74904, Qiagen) including an on-column DNase treatment (69104, Qiagen).

RNA-seq library construction was performed with the TruSeq kit (Illumina) and 150-bp paired-end sequencing with BGISEQ-500 was performed by BGI Tech Solutions (Hong Kong). Obtained raw fastq reads were quality controlled using FastQC (https://www.bioinformatics.babraham.ac.uk/projects/fastqc/). Cleaned reads were mapped to the *M. polymorpha* genome (v6.1, https:://marchantia.info/) using Hisat2 for paired-end reads (47), followed by counting reads per gene with FeatureCounts using default parameters (48). Finally, genes with low counts (<90 across all samples together) were removed and differentially expressed genes were calculated with DESeq2 (49).

### DNA-binding residues identification

DNA-binding residues were identified on the previously solved structures of AtARF1 DBD (PDB: 6YCQ) (14). Direct and indirect interactions were identified in PyMOL Molecular Graphics System (version 2.5.5, Schrödinger, LLC) between ARF and DNA molecules with a cutoff of 3.2 Å for hydrogen bonds, hydrophobic interactions (i.e., P184 and T1) by VdW radii, and π-interactions using the *distance* command in mode 5 with a cutoff of 6.5 Å.

### Structure conservation analyses

Conservation score values for ARF DBD residues were calculated using the ConSurf webserver (29). AtARF1 (6ycq, chain A) was used as query for the HMMER search (1 iteration), 0.0001 E-value cutoff, UNIREF-90 database). Conservation analysis was performed using default settings, except that homologues were filtered between 35-95% identity with AtARF1. Average conservation scores and standard deviations for specific domains were calculated by subsetting each domain and extracting conservation scores. The relative accessibility (sasa) of a residue – 0-30% accessible for buried, 31-100% for accessible residues – was used to identify buried (0-30% sasa) and accessible (31-100% sasa) residues.

### Protein structure prediction

Structures of ARFs were predicted using AlphaFold2 (Jumper *et al*., 2021) with default settings. Full-length ARF sequences were used for structure prediction, and middle region and PB1 manually trimmed to cover the DBD or B3. Structures were aligned in PyMOL using the *align* command with 5 cycles of outlier rejection.

### Molecular Docking and structural analysis

Molecular docking of DNA-binding variants was performed using HADDOCK (50). The MpARF2 structure (PDB: 6SDG) was used as template. The MpARF2 dimer (chain A and B) and DNA (Chain C and D) were saved as separate .pdb files. Chains B and D were renumbered (B +400, D +30) to conform with the unique identifier requirement of HADDOCK. Mutants in DNA-binding residues were made in both chains using the *mutation* tool of PyMOL followed by energy minimization of the entire dimer in Swiss PDB Viewer (51).

HADDOCK was performed in ‘easy’ mode with default settings. Active DNA residues were: DNA: 1, 2, 3, 4, 5, 6, 14, 15, 16, 17, 18, 19, 31, 32, 33, 34, 35, 36, 44, 45, 46, 47, 48, 49; MpARF2: 136, 141, 145, 148, 186, 189, 191, 196, 199, 536, 541, 545, 548, 586, 589, 591, 596, 599. No passive residues for the DNA, but for MpARF2: 131, 134, 135, 140, 149, 187, 188, 531, 534, 535, 540, 549, 587, 588. After a HADDOCK run, four structures per cluster were used and aligned to the original MpARF2 structure in PyMOL. Since we assumed our MpARF2 mutations would cause minimal changes to the DNA structure, we selected the cluster with the least RMSD compared with the original (∼0.5 RMSD averaged for all structures shown in this chapter). Representative HADDOCK structures were shown in the figures. All structural figures were made with PyMOL.

## Acknowledgements

We thank team members Sumanth Mutte and Shubhajit Das for support with the RNA-seq, and Erik van der Velde, Iraes Rabbers, Yiyun Li and Simone van Roosmalen for technical support. This work was supported by funding from the Nederlandse Organisatie voor Wetenschappelijk Onderzoek (NWO grant no. GSGT.GSGT.2018.013 to J.R. and OCENW.KLEIN.027 to D.W.) by a Marie Skłodowska-Curie Individual Fellowship (H2020-MSCA-IF-2020) to J.H.-G. and a Research Grant from the Human Frontiers Research Program (grant RGP0015/2022 to D.W.).

## References

1. J. L. Bowman, E. Flores Sandoval, H. Kato, On the Evolutionary Origins of Land Plant Auxin Biology. Cold Spring Harb Perspect Biol 13 (2021).

2. S. B. Li, Z. Z. Xie, C. G. Hu, J. Z. Zhang, A Review of Auxin Response Factors (ARFs) in Plants. Front Plant Sci 7, 47 (2016).

3. D. Weijers, D. Wagner (2016) Transcriptional Responses to the Auxin Hormone. in Annual Review of Plant Biology, pp 539–574.

4. H. Suzuki, H. Kato, M. Iwano, R. Nishihama, T. Kohchi, Auxin signaling is essential for organogenesis but not for cell survival in the liverwort Marchantia polymorpha. Plant Cell 35, 1058–1075 (2023).

5. X. Tan et al., Mechanism of auxin perception by the TIR1 ubiquitin ligase. Nature 446, 640 (2007).

6. S. B. Tiwari, AUX/IAA Proteins Are Active Repressors, and Their Stability and Activity Are Modulated by Auxin. The Plant Cell Online 13, 2809–2822 (2001).

7. T. Ulmasov, J. Murfett, G. Hagen, T. J. Guilfoyle, Aux/IAA proteins repress expression of reporter genes containing natural and highly active synthetic auxin response elements. Plant Cell 9, 1963–1971 (1997).

8. T. Ulmasov, G. Hagen, T. J. Guilfoyle, ARF1, a transcription factor that binds to auxin response elements. Science 276, 1865–1868 (1997).

9. T. Ulmasov, Z. B. Liu, G. Hagen, T. J. Guilfoyle, Composite structure of auxin response elements. Plant Cell 7, 1611–1623 (1995).

10. D. R. Boer et al., Structural basis for DNA binding specificity by the auxin-dependent ARF transcription factors. Cell 156, 577–589 (2014).

11. M. H. Nanao et al., Structural basis for oligomerization of auxin transcriptional regulators. Nat Commun 5, 3617 (2014).

12. M. Fontana et al., Cooperative action of separate interaction domains promotes high-affinity DNA binding of Arabidopsis thaliana ARF transcription factors. Proc Natl Acad Sci U S A 120, e2219916120 (2023).

13. T. Ulmasov, G. Hagen, T. J. Guilfoyle, Activation and repression of transcription by auxin-response factors. Proc Natl Acad Sci U S A 96, 5844–5849 (1999).

14. A. Freire-Rios et al., Architecture of DNA elements mediating ARF transcription factor binding and auxin-responsive gene expression in Arabidopsis. Proc Natl Acad Sci U S A 117, 24557–24566 (2020).

15. E. V. Zemlyanskaya, D. S. Wiebe, N. A. Omelyanchuk, V. G. Levitsky, V. V. Mironova, Meta-analysis of transcriptome data identified TGTCNN motif variants associated with the response to plant hormone auxin in Arabidopsis thaliana L. J Bioinform Comput Biol 14, 1641009 (2016).

16. P. Cherenkov et al., Diversity of cis-regulatory elements associated with auxin response in Arabidopsis thaliana. J. Exp. Bot. 69, 329–339 (2018).

17. R.C. O’Malley et al., Cistrome and Epicistrome Features Shape the Regulatory DNA Landscape. Cell 165, 1280–1292 (2016).

18. M. Galli et al., The DNA binding landscape of the maize AUXIN RESPONSE FACTOR family. Nat Commun 9, 4526 (2018).

19. A. Stigliani et al., Capturing Auxin Response Factors Syntax Using DNA Binding Models. Mol Plant 12, 822–832 (2019).

20. S. K. Mutte et al., Origin and evolution of the nuclear auxin response system. Elife 7 (2018).

21. R. Martin-Arevalillo et al., Evolution of the Auxin Response Factors from charophyte ancestors. PLoS Genet. 15, e1008400 (2019).

22. J. Hernández-García et al., Evolutionary Origins and Functional Diversification of Auxin Response Factors. bioRxiv 10.1101/2024.08.14.607941, 2024.2008.2014.607941 (2024).

23. H. Kato et al., Auxin-Mediated Transcriptional System with a Minimal Set of Components Is Critical for Morphogenesis through the Life Cycle in Marchantia polymorpha. PLoS Genet. 11, e1005084 (2015).

24. H. Kato et al., Design principles of a minimal auxin response system. Nat Plants 6, 473–482 (2020).

25. R. Martin-Arevalillo et al., Structure of the Arabidopsis TOPLESS corepressor provides insight into the evolution of transcriptional repression. Proc Natl Acad Sci U S A 114, 8107–8112 (2017).

26. M. van Dijk, K. M. Visscher, P. L. Kastritis, A. M. Bonvin, Solvated protein-DNA docking using HADDOCK. J. Biomol. NMR 56, 51–63 (2013).

27. D. Kosztin, T. C. Bishop, K. Schulten, Binding of the estrogen receptor to DNA. The role of waters. Biophys. J. 73, 557–570 (1997).

28. C. A. Davey, D. F. Sargent, K. Luger, A. W. Maeder, T. J. Richmond, Solvent mediated interactions in the structure of the nucleosome core particle at 1.9 a resolution. J. Mol. Biol. 319, 1097–1113 (2002).

29. H. Ashkenazy et al., ConSurf 2016: an improved methodology to estimate and visualize evolutionary conservation in macromolecules. Nucleic Acids Res. 44, W344–350 (2016).

30. J. Jumper et al., Highly accurate protein structure prediction with AlphaFold. Nature 596, 583–589 (2021).

31. M. Varadi et al., AlphaFold Protein Structure Database: massively expanding the structural coverage of protein-sequence space with high-accuracy models. Nucleic Acids Res. 50, D439–D444 (2021).

32. C. Beaulieu et al., The Marchantia pangenome reveals ancient mechanisms of plant adaptation to the environment. bioRxiv 10.1101/2023.10.27.564390 (2023).

33. E. H. Rademacher et al., Different auxin response machineries control distinct cell fates in the early plant embryo. Dev. Cell 22, 211–222 (2012).

34. E. H. Rademacher et al., A cellular expression map of the Arabidopsis AUXIN RESPONSE FACTOR gene family. Plant J. 68, 597–606 (2011).

35. Y. Okushima, I. Mitina, H. L. Quach, A. Theologis, AUXIN RESPONSE FACTOR 2 (ARF2): a pleiotropic developmental regulator. Plant J. 43, 29–46 (2005).

36. Y. Okushima et al., Functional genomic analysis of the AUXIN RESPONSE FACTOR gene family members in Arabidopsis thaliana: unique and overlapping functions of ARF7 and ARF19. Plant Cell 17, 444–463 (2005).

37. D. W. Anderson, A. N. McKeown, J. W. Thornton, Intermolecular epistasis shaped the function and evolution of an ancient transcription factor and its DNA binding sites. eLife 4, e07864 (2015).

38. T. N. Starr, L. K. Picton, J. W. Thornton, Alternative evolutionary histories in the sequence space of an ancient protein. Nature 549, 409–413 (2017).

39. N. Butel, Y. Qiu, W. Xu, J. Santos-Gonzalez, C. Kohler, Parental conflict driven regulation of endosperm cellularization by a family of Auxin Response Factors. Nat Plants 10, 1018–1026 (2024).

40. C. Shen et al., Functional analysis of the structural domain of ARF proteins in rice (Oryza sativa L.). J. Exp. Bot. 61, 3971–3981 (2010).

41. S. S. Sugano et al., CRISPR/Cas9-mediated targeted mutagenesis in the liverwort Marchantia polymorpha L. Plant Cell Physiol. 55, 475–481 (2014).

42. H. Kato et al., The Roles of the Sole Activator-Type Auxin Response Factor in Pattern Formation of Marchantia polymorpha. Plant Cell Physiol. 58, 1642–1651 (2017).

43. T. Gera, F. Jonas, R. More, N. Barkai, Evolution of binding preferences among whole-genome duplicated transcription factors. Elife 11 (2022).

44. J. C. Perez et al., How duplicated transcription regulators can diversify to govern the expression of nonoverlapping sets of genes. Genes Dev. 28, 1272–1277 (2014).

45. C. Sayou, M. Monniaux, M. Nanao, E. Moyroud, A promiscuous intermediate underlies the evolution of LEAFY DNA binding specificty. Science 343, 645–648 (2014).

46. S. Berg et al., ilastik: interactive machine learning for (bio)image analysis. Nat. Methods 16, 1226–1232 (2019).

47. D. Kim, J. M. Paggi, C. Park, C. Bennett, S. L. Salzberg, Graph-based genome alignment and genotyping with HISAT2 and HISAT-genotype. Nat. Biotechnol. 37, 907–915 (2019).

48. Y. Liao, G. K. Smyth, W. Shi, featureCounts: an efficient general purpose program for assigning sequence reads to genomic features. Bioinformatics 30, 923–930 (2014).

49. M. I. Love, W. Huber, S. Anders, Moderated estimation of fold change and dispersion for RNA-seq data with DESeq2. Genome Biology 15, 550 (2014).

50. G. C. P. van Zundert et al., The HADDOCK2.2 Web Server: User-Friendly Integrative Modeling of Biomolecular Complexes. J. Mol. Biol. 428, 720–725 (2016).

51. N. Guex, M. C. Peitsch, SWISS-MODEL and the Swiss-PdbViewer: an environment for comparative protein modeling. Electrophoresis 18, 2714–2723 (1997).

