## Supplementary Figures 1-10 for "An ultra-conserved ARF-DNA interface underlies auxin-triggered transcriptional response"

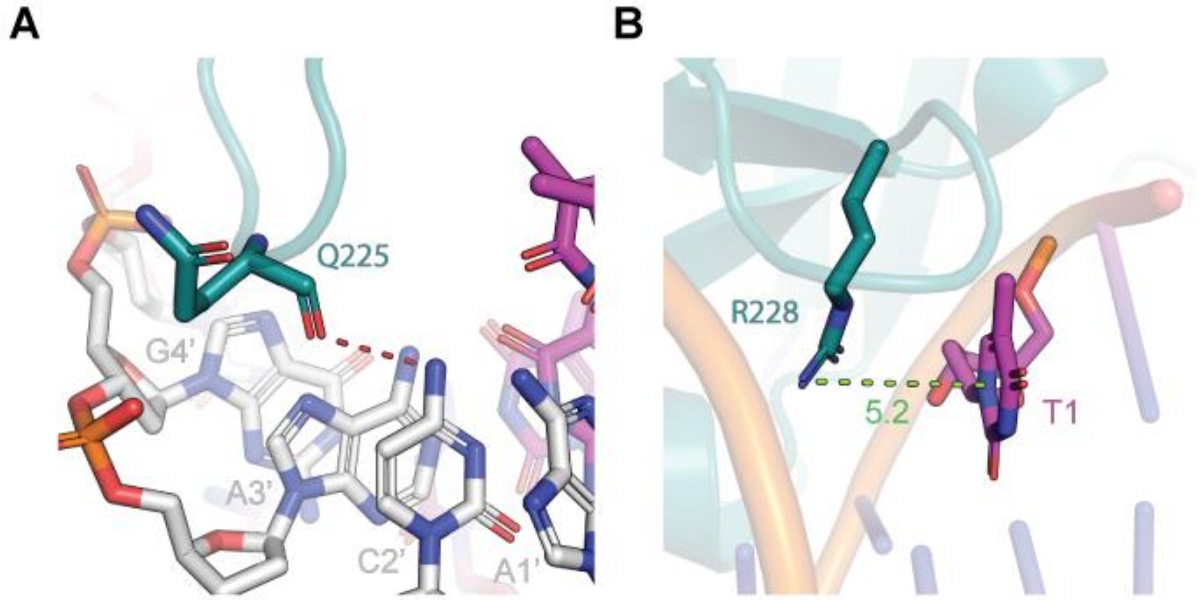


**Figure S1 | DNA-binding residues in MpARF2. (A)** The MpARF2-Q225 forms hydrogen bonds with C2' using the carboxyl-group similar to AtARF1-Q183. **(B)** MpARF2-R228 forms pi-interactions with T1. (A & B) Interacting residues of MpARF2 (teal) and DNA (white/magenta) represented as sticks. Dashed lines indicate hydrogen bonds (red) or pi-interactions (green), the latter with distance in Å

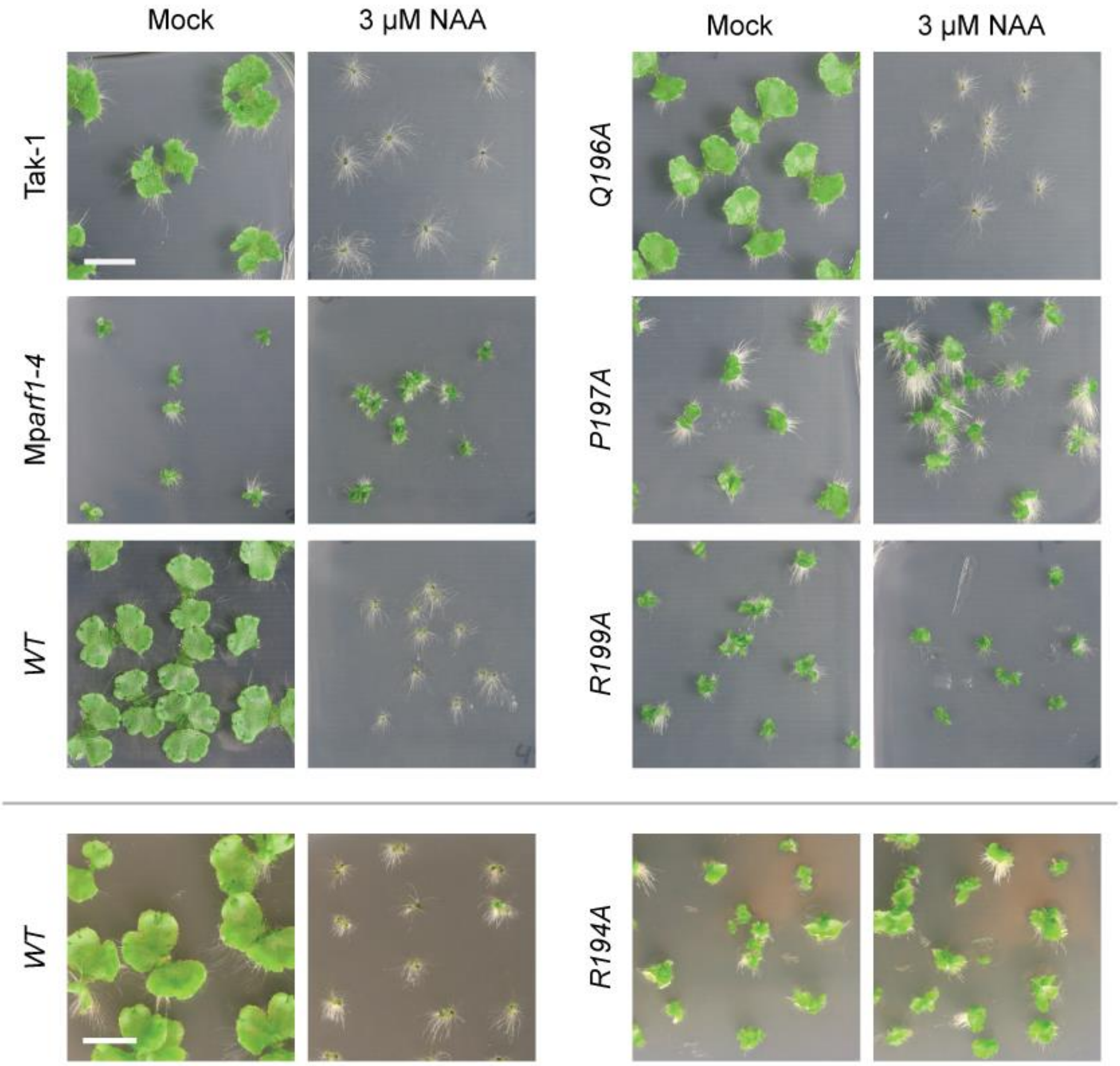


**Figure S2 | Pictures of *M. polymorpha* plants with alanine variants of MpARF1.** Pictures of 14 day old plants of Tak-1, Mp*arf1-4*, and Mp*arf1-4* complemented with either wild type MpARF1 (WT) or alanine mutation variants, grown on medium without (mock) or with 3 μM NAA. Pictures above and below the horizontal line correspond to different experiments. Scale bar is 1 cm

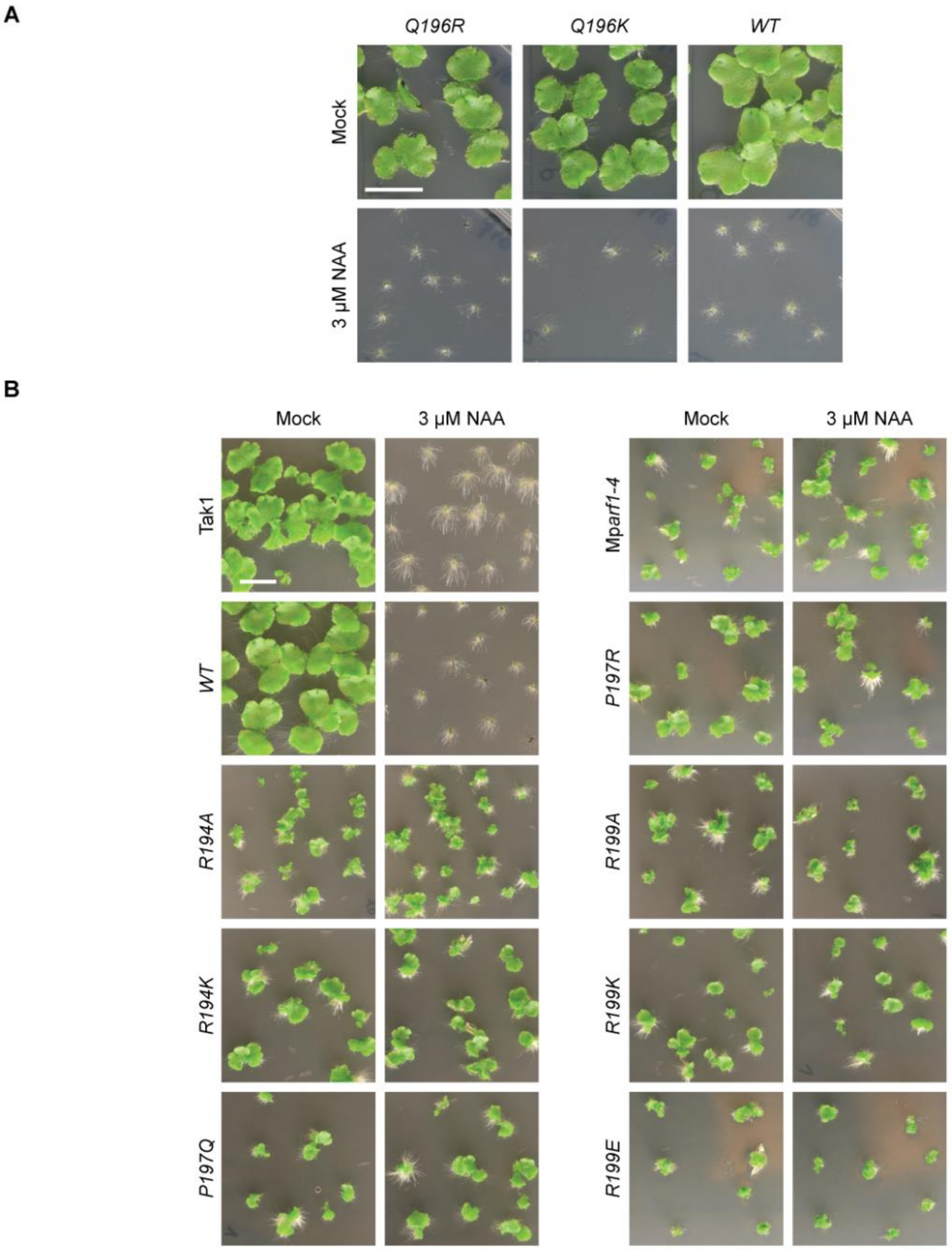


**Figure S3 | Pictures of *M. polymorpha* plants with MpARF1 variants based on variation observed in land plant ARFs. (A-B)** Pictures of 14-day-old plants of Tak-1, Mp*arf1-4*, and Mp*arf1-4* transformed with MpARF1 constructs without mutations (WT) or with indicated mutations. Between 3 and 8 independent lines were screened, representative lines are shown. A and B are independent experiments. Scale bar is 1 cm.

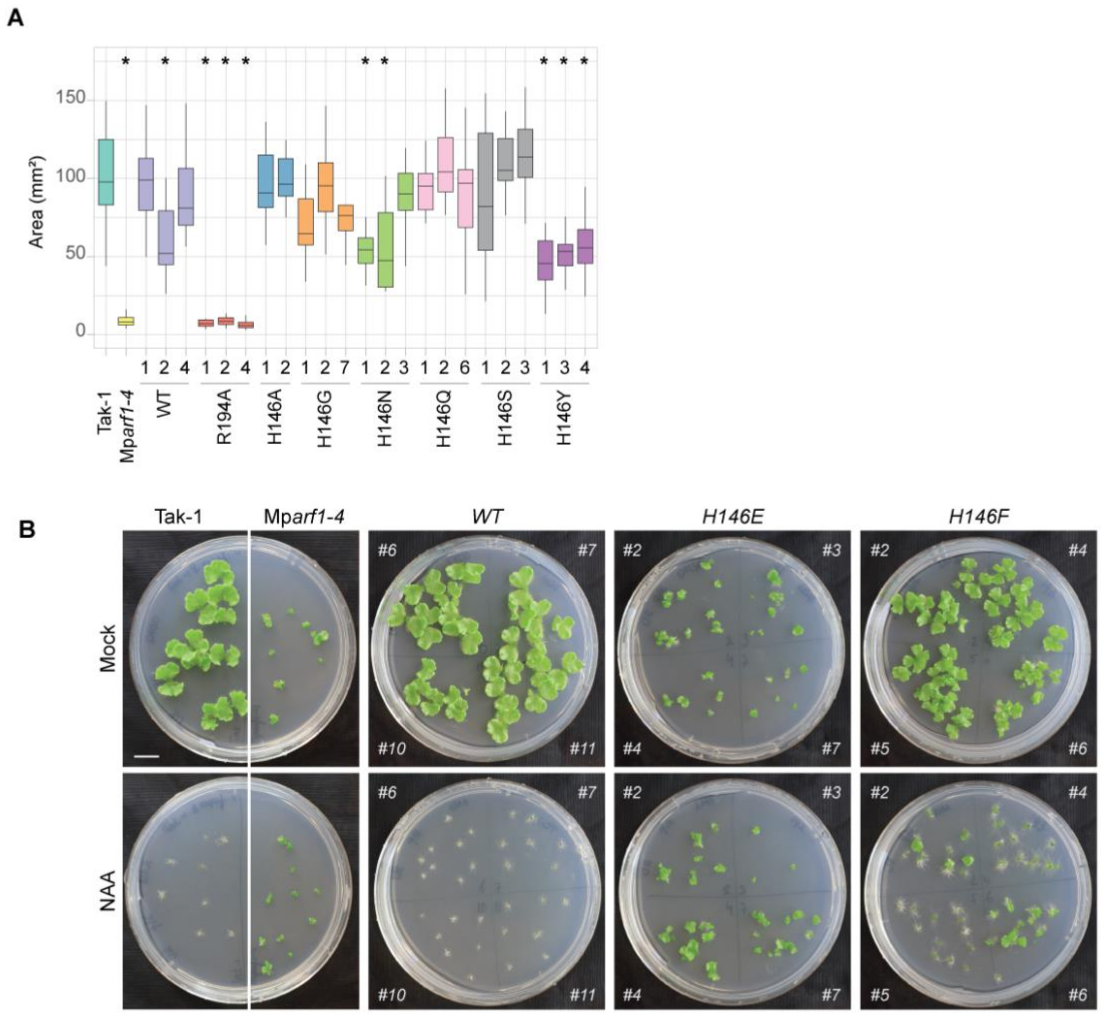


**Figure S4 | Quantification and pictures of *M. polymorpha* plants with MpARF1-H146 variants based on variation observed in land plant ARFs. (A)** Thallus area ($mm^2$) of 14 day old plants of MpARF1 variants grown under standard conditions. Between 2 and 3 independent lines were screened, with 4 replicates per line (*n* = 4). **(B)** Pictures of 14 day old plants of wild type MpARF1 (WT) and MpARF1-H146E and H146F variants, grown on medium without (mock) or with 3 μM NAA. Seven independent lines were screened. Scale bar is 1 cm.

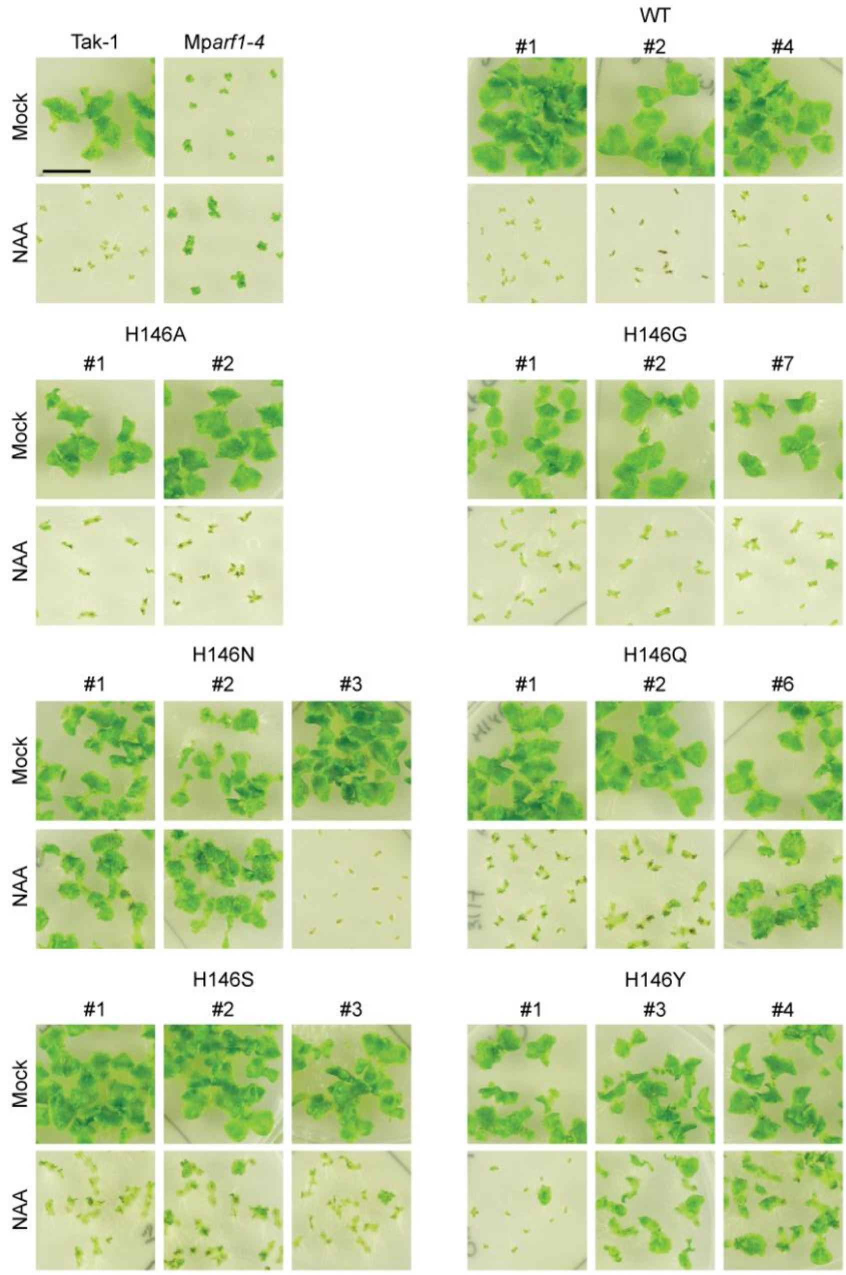


**Fig. S5 | Auxin response of MpARF1 variants.** Pictures of 14-day old plants of MpARF1 variants grown without (mock) or with 3 µM NAA. Indicated are genotype and the number of the individual line. Scale bar is 1 cm.

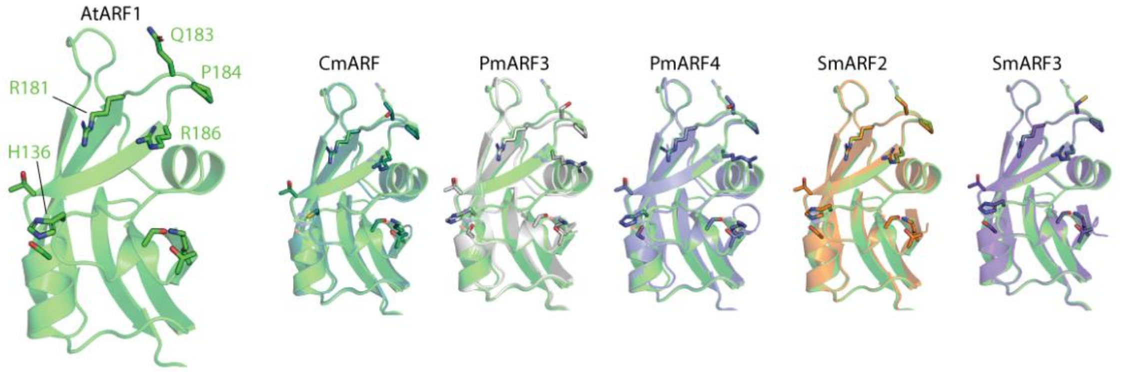


**Fig. S6 | Structural alignment of B3 domain and DNA-binding residues of AtARF1 and algal ARFs.** Left. The B3 domain of the AtARF1 crystal structure (PDB: 4ldx). DNA-binding residues are shown as sticks, with the main DNA-binding residues highlighted by name. Right. Structural alignment of AtARF1 with the predicted structures of varying algal ARFs, with homologous DNA-binding residues shown as sticks.

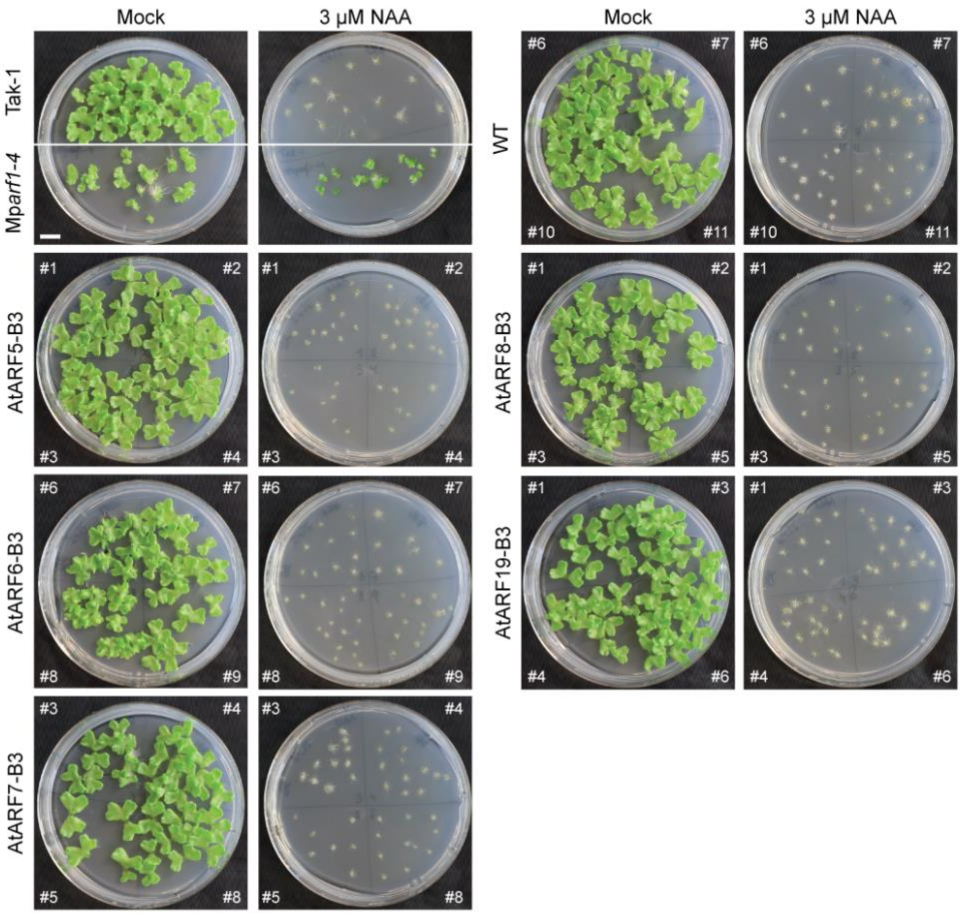


**Fig. S7 | Phenotype of plants with B3 swaps with AtARFs.** Pictures of 14-day old plants of wild type MpARF1 (WT) and B3 swap variants wherein the MpARF1-B3 was swapped for the B3 domain of AtARFs grown without (mock) or with 3 µM NAA. Between 4-8 independent lines were screened for each construct. (A) For each picture, the MpARF1 construct is written on the left, the white numbers inside the picture indicate the independent line.

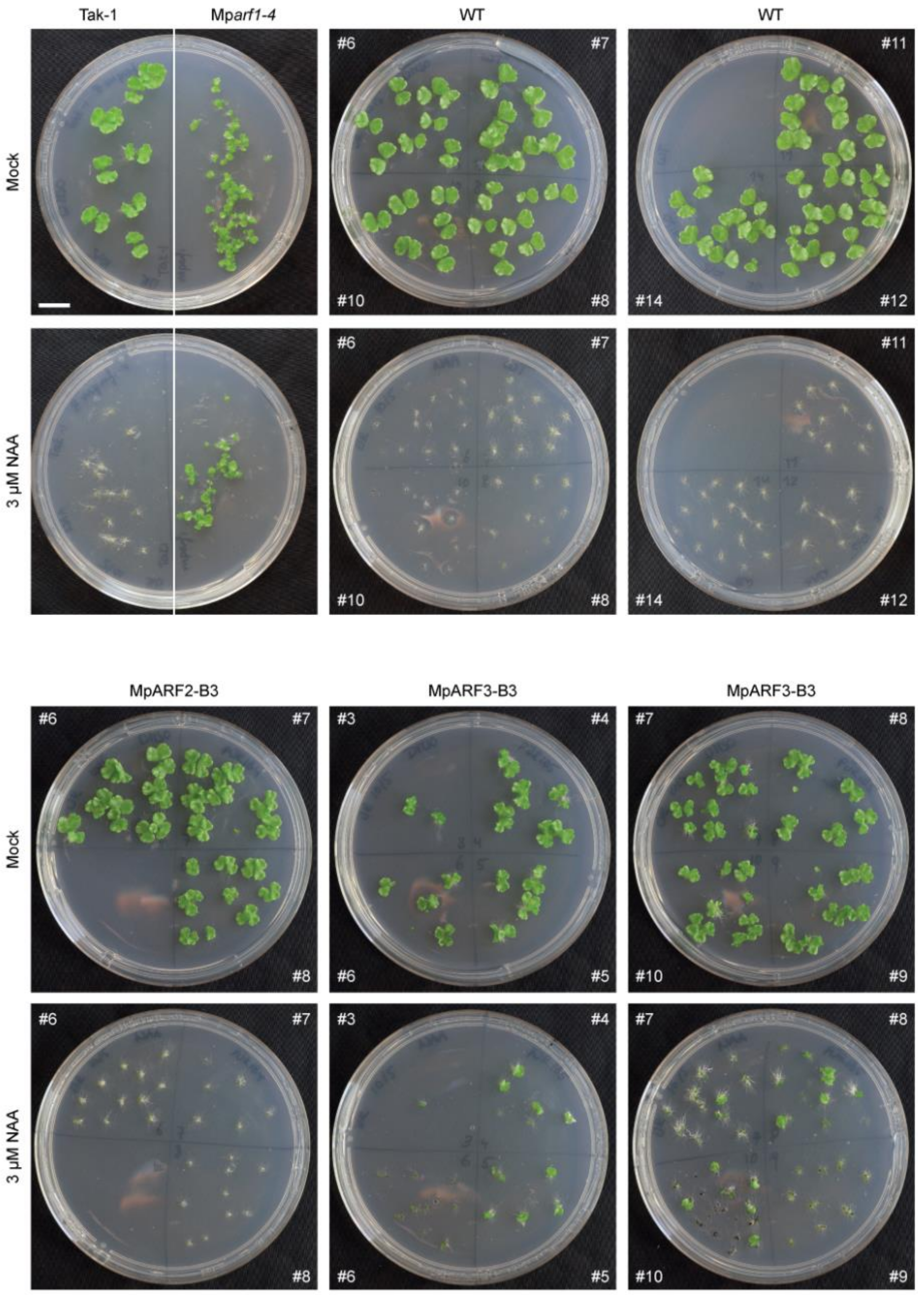


**Fig. S8 | Phenotype of plants with B3 swaps with MpARFs.** Pictures of 14-day old plants of wild type MpARF1 (WT) and B3 swap variants wherein the MpARF1-B3 was swapped for the B3 domain of MpARF2 or MpARF3 grown without (mock) or with 3 µM NAA. Between 4-8 independent lines were screened for each construct. (A) For each picture, the MpARF1 construct is written on the left, the white numbers inside the picture indicate the independent line.

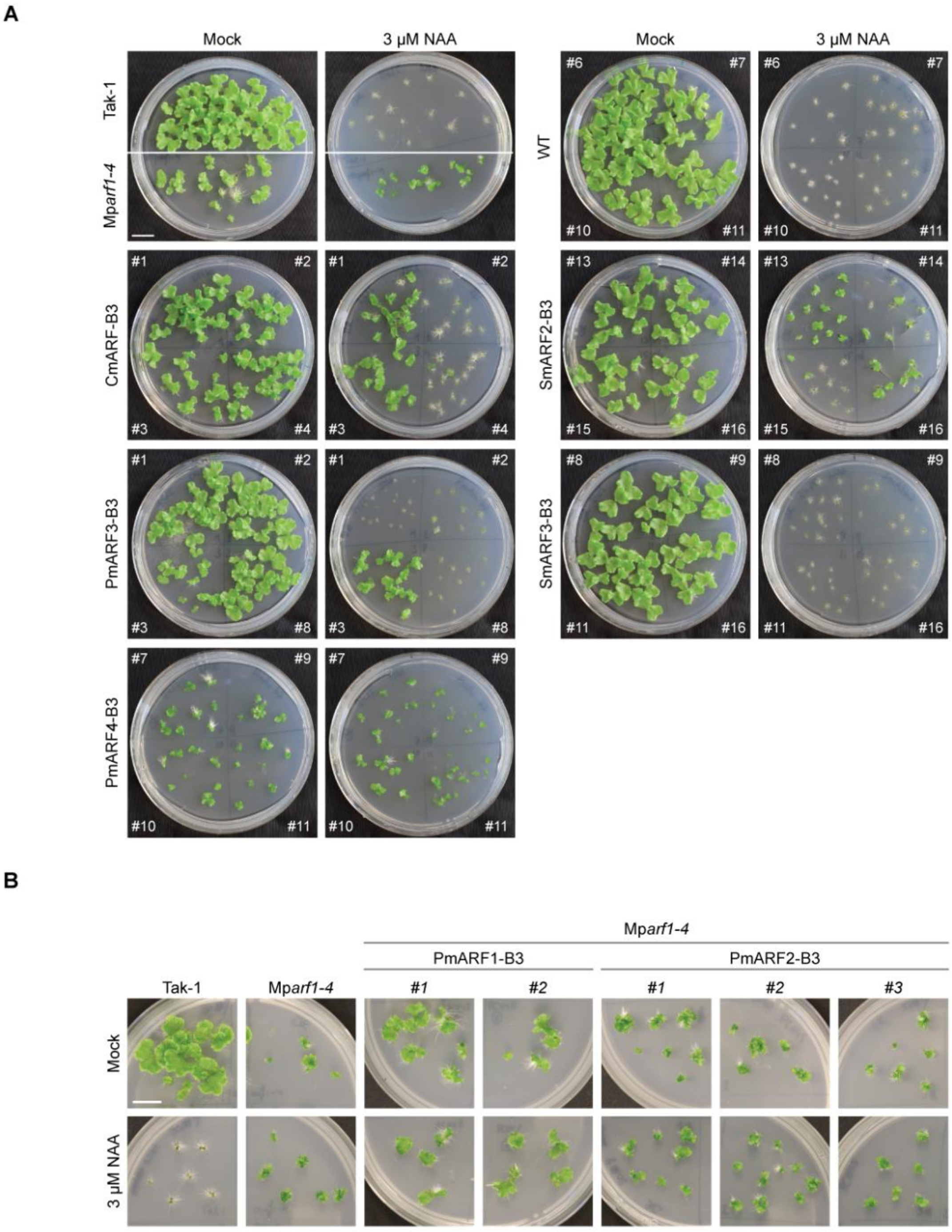


**Fig. S9 | Phenotype of plants with algal B3 swaps. (A & B)** Pictures of 14-day old plants of wild type MpARF1 (WT) and B3 swap variants wherein the MpARF1-B3 was swapped for the B3 domain of algal ARFs grown without (mock) or with 3 µM NAA. Between 3-8 independent lines were screened for each construct. (A) For each picture, the MpARF1 construct is written on the left, the white numbers inside the picture indicate the independent line. (B) For each picture, the construct and independent lines are indicated on the top. Scale bar is 1 cm.

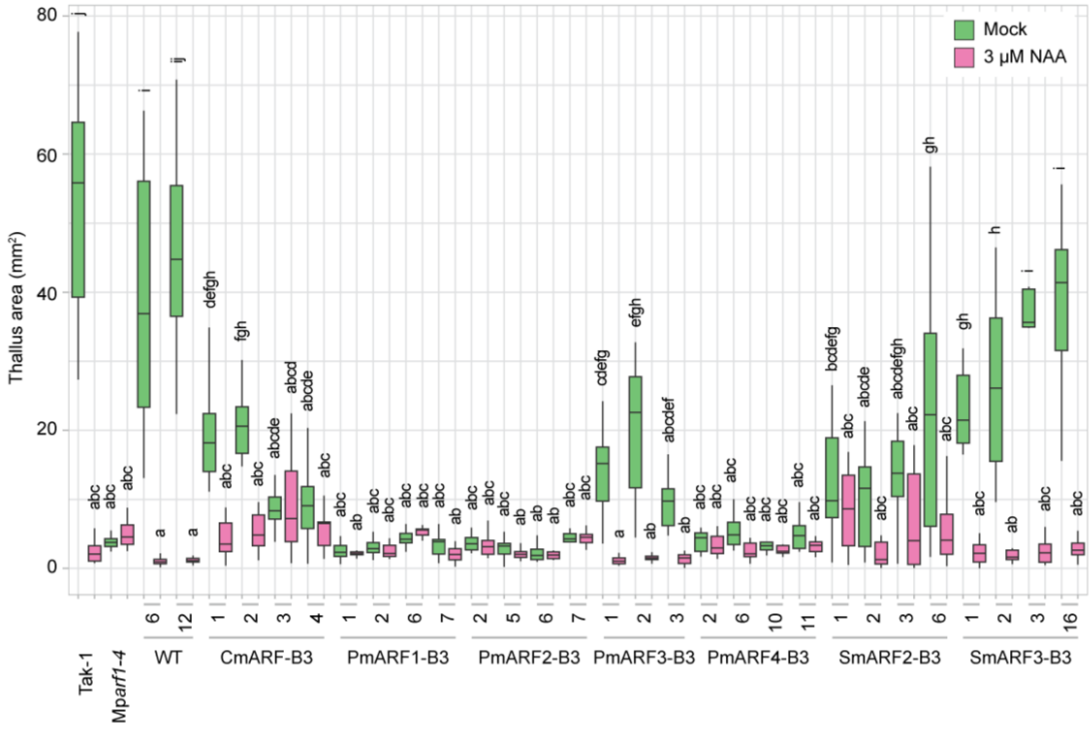


**Fig. S10 | Thallus area (mm$^2$) of 14-day old plants of MpARF1 with B3 swaps from algal ARFs.** Constructs and independent line numbers are indicated below the bar chart. Six plants ($n$ = 6) per line were quantified after 14 days grown on medium without (mock, green) or with 3 µM NAA (magenta). Lower caps letters in the graph indicate different statistical groups (ANOVA, posthoc test with Sidak correction, $p \leq 0.05$). Experiments for this graph were independently from Fig. S9.
